# Development and social dynamics of stone tool use in wild white-faced capuchin monkeys

**DOI:** 10.1101/2025.04.08.647785

**Authors:** Zoë Goldsborough, Meredith K.W. Carlson, Leonie S. Reetz, Margaret C. Crofoot, Brendan J. Barrett

## Abstract

Percussive tool use for extractive foraging allows animals to access otherwise inaccessible resources and forage more efficiently, with potentially important implications for their fitness. The development of tool use proficiency has been well-documented in nut-cracking chimpanzees and robust capuchins, and in shellfish-cracking long-tailed macaques, where mothers and proficient tool users are the most important models for social learning. However, little is known about how tool use develops in populations where opportunities for social learning are scarce. White-faced capuchins (*Cebus capucinus imitator*) on Jicarón Island, Panama, provide a unique case to consider this question: stone tool use is entirely male-biased, meaning juveniles cannot learn from their mothers, and reduced group cohesion further limits social learning opportunities. Here, we investigate the acquisition and development of stone tool proficiency in this population using a year-long dataset from camera traps placed at two experimental anvils. We assess differences in proficiency between age classes, examine the development of tool use proficiency over time, and explore patterns of social attention during tool use. We show that juvenile capuchins are less proficient than subadults and adults, but their proficiency remains stable over the course of one year, suggesting that skill development may require prolonged practice or physical maturation. In contrast to other primates, social learning opportunities on Jicarón appear limited and scrounging is rare, yet when social attention occurs, we find robust patterns. Social attention to tool use mostly comes from juveniles too young to use tools themselves, who observe proficient subadults that tolerate scrounging. Our results contribute to the understanding of how complex tool use behaviours are acquired and maintained in primates. They highlight that even when tool use is largely solitary, directed social attention and social tolerance play an important role in the development of tool use proficiency.

**Highlights:**

- Juvenile white-faced capuchins are less proficient tool users than (sub)adults
- Development of capuchins’ tool use proficiency is a slow process
- Social learning opportunities on Jicarón island are limited and social attention rare
- Attention was selective, juveniles to proficient sub(adults) who tolerated scrounging
- Non-tool-using juveniles paid most attention, so likely important for acquisition

## Introduction

Tool use is a striking example of behavioural flexibility in animals, and the use of tools for extractive foraging can increase fitness by allowing faster access to a greater variety of resources (Biro et al., 2013). However, becoming proficient in tool use can take years, depending on the complexity of the behaviour (C. Boesch et al., 2019). A behaviour is considered tool use when an animal: i) uses an object which is not part of their own body, ii) which is unattached to the surrounding environment (or attached and manipulatable (Shumaker et al., 2011)), and iii) the animal manipulates the object to reach a useful outcome (Beck, 1980). To describe complexity of tool use behaviour, a distinction can be made between first-order tool use (involving a single object, e.g., pounding a nut on a fixed surface) and second-order tool use (involving two objects, e.g., pounding a nut on a fixed surface using a hammerstone) (Fragaszy et al., 2004). Second-order tool use is thought to reflect greater cognitive complexity, as it requires more planning and physical coordination to correctly bring together all separate objects to obtain the desired outcome. Percussive stone technology—particularly extractive foraging using hammerstone and anvil tool use—is the focus of substantial scientific interest. This is due to its high level of complexity and its relative rarity in the animal kingdom, despite its prominent (and archaeologically preservable) role in human evolution (de la Torre & Hirata, 2015). The only non-human wild primates that regularly use stone tools for extractive foraging are chimpanzees (*Pan troglodytes* (Sugiyama & Koman, 1979)), long-tailed macaques (*Macaca fascicularis* spp. (Gumert & Malaivijitnond, 2012; Muhammad et al., 2024)), robust capuchins (*Sapajus* spp. (Canale et al., 2009; Lima et al., 2024; Ottoni & Mannu, 2001; Spagnoletti et al., 2011)), and gracile capuchins (*Cebus* spp. (Araujo et al., 2021; Barrett et al., 2018)). Among *Macaca* and *Cebus*, tool use is further restricted to only a few groups. Examining how stone tool use is acquired and how proficiency develops is key to understanding how a tool use tradition is maintained within a group, as well as how it spreads (or does not spread) between groups.

For particularly complex techniques like percussive stone tool use, acquisition of the skill may require sufficient exposure at a young age. In chimpanzees, insufficient exposure to nut-cracking during a critical learning period (between 3-10 years) may hinder or completely impede learning of the behaviour in adulthood (Biro et al., 2003; Inoue-Nakamura & Matsuzawa, 1997; Matsuzawa, 1994). In robust capuchins (De Resende et al., 2008) and macaques (Tan, 2017), acquisition of tool use also mostly occurs early in life. Exposure to tool use has an important social component: in both chimpanzees and robust capuchins, immature individuals are thought to acquire nut-cracking via social learning processes such as local enhancement, emulation, and social facilitation (Biro et al., 2003; Coelho et al., 2015; Eshchar et al., 2016; Ottoni et al., 2005). Though social learning has been argued not to be critical to the acquisition of nut-cracking in captive chimpanzees (Neadle et al., 2020), evidence from wild chimpanzees continues to support its important role (Koops et al., 2022).

In chimpanzees, most transmission of tool use behaviours occurs from mothers to their offspring (C. Boesch, 1991; Lonsdorf, 2005). Juveniles spend a great deal of time with their mothers during the sensitive learning period, and mothers are highly tolerant of their offspring observing their tool use events and scrounging on opened nuts. There is also some evidence of female chimpanzees providing learning opportunities for their offspring through “scaffolding” (sharing nut-cracking materials such as hammers with their offspring) (Estienne et al., 2019). In contrast, robust capuchins mothers were not common models to their offspring. Rather, juveniles preferentially observed tool use events by proficient tool users (Ottoni et al., 2005), who were older and also more dominant than themselves (Coelho et al., 2015). Social closeness or physical proximity did not affect who juveniles chose to observe. However, they did prefer individuals who were more tolerant of observers, and scrounging occurred in 75% of tool use events (Coelho et al., 2015).

High inter-individual social tolerance is thought to be an important pre-condition for tool use acquisition (van Schaik et al., 1999), as observational learning of a complex skill requires close visual attention to the tool use activities. Being in such close proximity to an intolerant tool-user is risky for an observer, who risks being chased off or attacked. High social tolerance often provides the opportunity to scrounge, which not only gives naïve individuals the chance to observe the end-result of tool use, but also allows them a share of the payoff—thus increasing their motivation to acquire tool use themselves (Fragaszy et al., 2013). After acquisition, honing of tool use skill can be a lengthy process. Younger, active learners spend more time on acquiring tool use and are less proficient than experienced adults (Tan, 2017). However, in bearded capuchins (*Sapajus libidinosus*), juveniles were not found to make more mistakes than adults during tool use. Rather, juveniles’ lower proficiency mainly resulted from less expertise in how to grip the hammerstone (Fragaszy et al., 2020). Due to differences in skill and/or strength, younger individuals may also be limited in what resources they can open (Spagnoletti et al., 2011). Furthermore, percussive stone tool use might have an upper age limit. In chimpanzees, progressive old age led to a reduction in both nut-cracking frequency and efficiency (Howard-Spink et al., 2024).

While the ontogeny of stone tool use for nut-cracking is well studied in chimpanzees and robust capuchins, little is known about gracile capuchins, who have only recently been added to the list of non-human primate species which habitually using stone tools. In particular, the white-faced capuchins (*Cebus capucinus imitator*) living in Coiba National Park, Panamá, are an important study system for studying tool use acquisition and proficiency development, as their social learning opportunities seem to differ from those of other tool-using primates. White-faced capuchins on the islands of Coiba and Jicarón habitually use stone tools to crack open a variety of food items, including fruits like sea almonds (*Terminalia catappa*), coconuts (*Cocos nucifera*) and palm fruits (*Bactris major*, *Astrocaryum standleyanum*) as well as invertebrates like Halloween crabs (*Gecarcinus quadratus*), hermit crabs (*Coenobita compressus*), nerite snails (*Nerita sp.*) and other freshwater mollusks (Barrett et al., 2018; Monteza-Moreno, Dogandžić, et al., 2020). However, on Jicarón, despite this tradition persisting for at least 17 years, it seems entirely localized to a 1.5 km stretch of coast, occupied by 2-3 neighbouring capuchin groups (Goldsborough et al., 2023). Only one group of capuchins appears to use tools habitually on the full variety of resources, while their neighbours have only been observed to use tools in the intertidal zone or in stream beds. In the most frequent tool-using group, stone tool use is fully male-biased: only males have ever been observed to use stone tools, despite females regularly using tools on the neighbouring island of Coiba (Goldsborough et al., 2024). The absence of tool use by females means that juvenile capuchins on Jicarón cannot learn tool use from their mothers, as is common in chimpanzees. Furthermore, this group of tool-using capuchins appear to have reduced group cohesion (Goldsborough et al., 2025), further limiting social learning opportunities. Taken together, the tool-using capuchins on Jicarón present a unique situation where tool use is highly localized, yet persistent, and naive individuals only have limited opportunity to observe tool use due to both less social cohesion and the male-exclusive nature of the behaviour, which makes mothers unavailable as a model. Given these constraints, understanding how tool use is acquired and develops in this group provides insights into the flexibility of social and individual learning mechanisms in primates.

Herein, we provide a first exploration of the acquisition and development of stone tool use proficiency by one group of white-faced capuchins on Jicarón island. Using a year of data from camera traps on two experimental anvils, we compare features of tool use proficiency– efficiency and mistakes– between juvenile, subadult, and adult capuchins. Furthermore, we present a longitudinal investigation of the improvement in tool use skills by juveniles over a 1-year period, as well as an initial exploration of tool use skill in old age. Lastly, we consider potential mechanisms driving acquisition of stone tool use on Jicarón through examining social attention to tool use sequences. We explore who pays attention to tool use events, which tool users are most likely to receive social attention, and how social attention relates to scrounging and tool user proficiency. In doing so, we provide a first reference of stone tool use proficiency development in white-faced capuchins, which will allow for comparison to other stone-tool-using primate species and which lays the groundwork for future studies into the maintenance and spread of tool use behaviour in this population.

## Methods

### Site and subjects

Jicarón island (2002 ha) is part of Coiba National Park, an archipelago consisting of nine islands and 100 islets off the Pacific coast of the Veraguas Province, Panama. Jicarón island lies approximately 60 kilometres from the mainland, and is estimated to have been separated from the mainland for 14 000-18 000 years (Titcomb & O’Dea, 2020). Archaeological evidence suggests that Jicarón island was used by indigenous communities from at least 250 CE until shortly before European contact(Isaza & Vrba, 2010), and today it is uninhabited, with only occasional visits by scientists and locals. White-faced capuchins occur on the three largest islands in Coiba National Park (Barrett et al., 2018). Jicarón has depauperate mammalian communities and a total absence of mammalian predators (Ibáñez et al., 1997) and white-faced capuchins show increased terrestriality (Monteza-Moreno, Crofoot, et al., 2020) and live at high densities (Barrett et al., 2018; Milton & Mittermeier, 1977) compared to mainland populations. The most active tool-using group on Jicarón has been monitored using unbaited camera traps since 2017, and has been using tools since at least 2004 (Barrett et al., 2018). Stone tool use for extractive foraging has been documented to occur at three types of sites in this group, distinguishable through differences in debris accumulation (see (Goldsborough et al., 2023) for more details). For the current study, we focused on tool use at high accumulation sites, from here on out referred to as “anvils”, where capuchins habitually use tools and large amounts of debris accumulate over time.

In 2022, the capuchin group was estimated to be comprised of around 20-25 individuals (5-6 adult males, 5-6 adult females, 4-5 subadults, 7-10 juveniles). These estimates are based on identifiable individuals (Table 1) as well as the maximum number of capuchins of a specific age-sex class observed together in one sequence. Ages were estimated by visual assessment based off of body size, shape, and facial appearance in comparison to demographically known mainland populations. We were unable to obtain precise estimates of group size due to the nature of data collection via camera trapping, where the visual field is always limited, and juveniles are difficult to identify reliably from images alone.

**Table 1:**
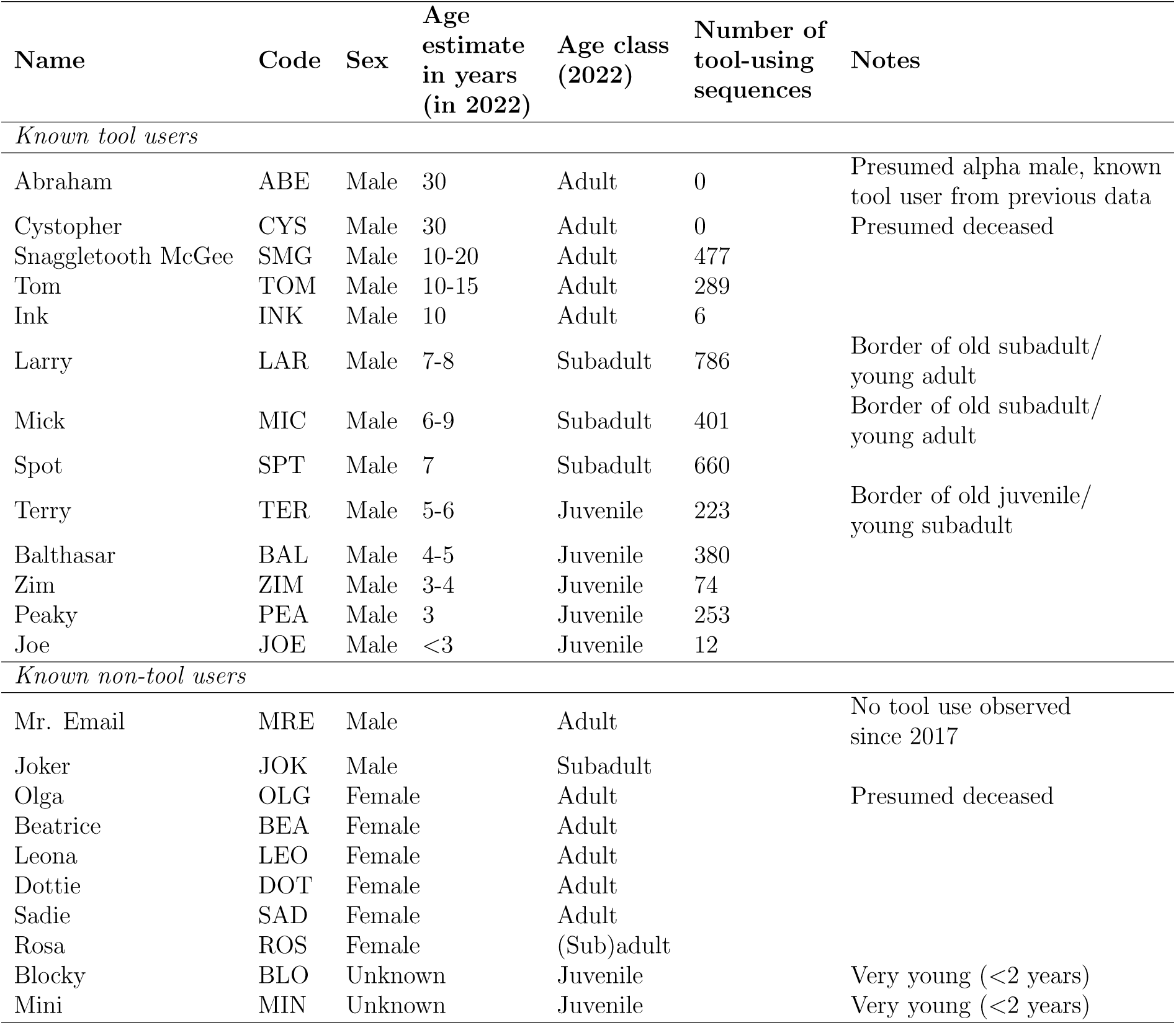
Overview of known individuals in the tool-using group, divided into known tool-users and known non-tool-users. Age estimates in years for tool users are rough approximations based on size and growth from 2017-2025. The borders for age classes are juveniles <6 years, subadults 6-8 years of age and adults *>*8 years.

### Experimental anvil set-up

We set up two experimental anvil sites in the range of the tool-using group in January 2022: a wooden anvil at a site which has been heavily used since 2017 (location name CEBUS-02), and a stone anvil at an entirely novel location under sea almond trees (*Terminalia catappa*) where there was no evidence of habitual tool use (EXP-ANV-01). To set up each experimental anvil, we first cleared the area of debris and placed a 1.8 m^2^ piece of nylon mesh on the ground, on which we placed the anvil. Anvils were selected from nearby based on the presence of a flat processing surface, as well as similarity in size to the preexisting anvil at CEBUS-02, which washed away prior to these experiments. Additionally, at each anvil, we supplied four uniquely marked hammerstones of varying sizes and weights sourced from the nearby intertidal zone (Figure 1A). While both anvils were placed under sea almond trees, no nuts were included in the experimental set-up. This setup was also designed to accommodate a concurrent archaeological study examining hammerstone selectivity and related tool-use patterns.

**Figure 1:**
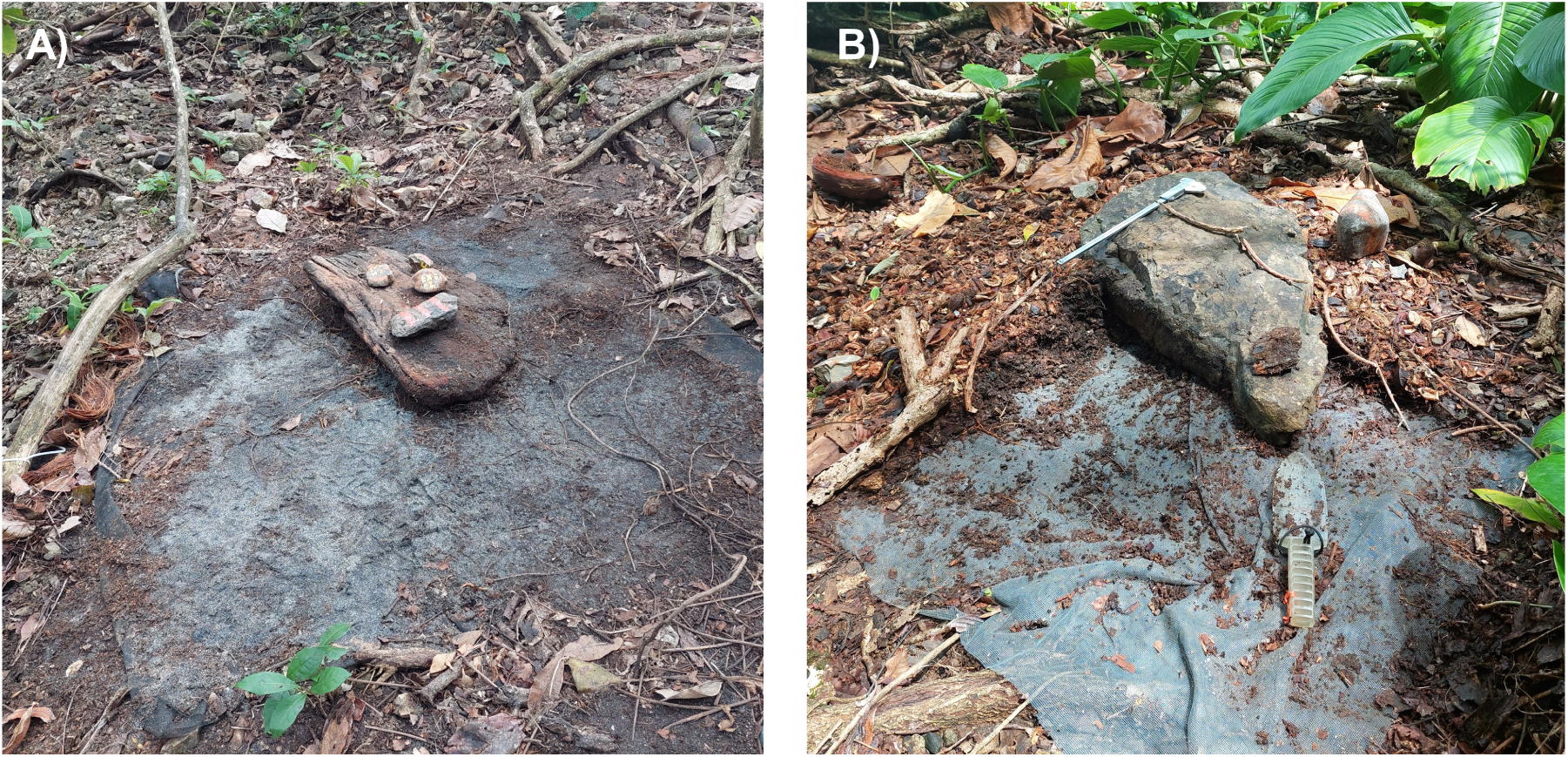
Photographs of the experimental anvils, showing. A) placement of CEBUS-02 in January 2022, and B) EXP-ANV-01 during excavation in July 2022. Photo credits belong to Brendan J. Barrett and Meredith Carlson.

Two camera traps were attached to trees and aimed at each experimental anvil: a close-up video camera (Reconyx UltraFire XP9) recording a 60 second video per trigger, and a wider-angle still (Reconyx Hyperfire HF2X) camera trap recording 30 images per trigger event without any between-trigger delays. Camera traps were placed in two deployments lasting approximately 6 months, and ran from January 2022 until January 2023. When camera traps were replaced in July 2022, both experimental anvils were excavated (Figure 1B), meaning all debris (food remains and tools) was collected down to the underlying mesh. Hammerstones and fragments found within the mesh perimeter were recovered and identified on the basis of markings and refits, when necessary. The remaining intact hammerstones were provided on the anvil during the second deployment.

### Coding

All videos containing tool use events were coded frame-by-frame in BORIS v. 8.21.8 (Friard & Gamba, 2016) by two coders (ZG and LR) in a split-half design. One coder coded days 1-15 and the other days 16-end of each month, alternating which coder coded the first versus second half across deployments. We considered individual tool use sequences, where the start is defined as the moment an individual placed an item on the anvil for processing and grabbed the hammerstone. A sequence ended when either: i) the item was opened and consumed ii) the individual relocated with the item to a place out of view of the camera or iii) the item was abandoned, which includes when the item was opened but not eaten. Per tool use sequence, we identified the age class (juvenile, subadult, or adult) of the tool user, and their identity when possible. We classified age based on visual appearance, with juveniles being individuals estimated to be *<*6 years old, subadults 6-8 years old, and adults *>*8 years old. We subsequently coded all behaviours by the tool user following an ethogram we established (Table S1, see also video ethogram https://keeper.mpdl.mpg.de/d/0c1b9853f3f342d8b3da/), coding features of tool use efficiency (e.g., number of successful pounds, number of misstrikes) and technique (e.g., type of pound [crouching, standing, jumping]). We also coded asocial (e.g., type of item processed, hammerstone material) and social factors (e.g., displacements, social attention, scrounging) on the level of the sequence (Table S2). A single video could contain one or more tool use sequences, and sequences could also span several videos if the individual was not finished processing when the video ended, and continued in the next triggered video. LR was trained by ZG on a sample set of data before coding independently. Inter-rater reliability was established by double-coding a randomly selected sample of 121 videos containing 203 sequences, with agreement assessed for all categorical, numerical, and timing variables (Table 2).

**Table 2:** Inter-rater reliability between two coders of all coded variables used in analyses. Categorical variables are summarised with Cohen’s k, numerical variables with ICC, and timing variables with mean absolute difference, SD, and proportion agreeing within 1 second

| ***Categorical variables*** |  |  |  |  |
| --- | --- | --- | --- | --- |
| **Variable** | **Cohen’s k** | **95% CI Lower** | **95% CI Upper** | **N Sequences** |
| Sequence outcome (e.g., opened) | 0.82 | 0.71 | 0.94 | 203 |
| Displacement | 0.82 | 0.74 | 0.90 | 203 |
| Social attention | 0.81 | 0.72 | 0.89 | 203 |
| Scrounging | 0.82 | 0.74 | 0.91 | 203 |
| Type of item (no colour of almendra) | 0.83 | 0.60 | 1.00 | 196 |
| colour of almendra (if both assigned) | 0.70 | 0.53 | 0.87 | 127 |
| Age of tool user | 0.98 | 0.96 | 1.00 | 203 |
| Identity of tool user (if both assigned) | 0.96 | 0.93 | 0.99 | 180 |
| ***Numerical variables*** |  |  |  |  |
| **Variable** | **ICC** | **95% CI Lower** | **95% CI Upper** | **N Sequences** |
| Number of pounds | 0.98 | 0.98 | 0.99 | 203 |
| Number of item repositions | 0.90 | 0.87 | 0.93 | 203 |
| Number of peels | 0.94 | 0.93 | 0.96 | 203 |
| Number of misstrikes | 0.73 | 0.65 | 0.78 | 203 |
| Number of item flies | 0.68 | 0.60 | 0.75 | 203 |
| Duration of sequence (s) | 0.99 | 0.99 | 0.99 | 203 |
| ***Timing variables*** |  |  |  |  |
| **Variable** | **Mean abs. diff.** | **SD abs. diff.** | **Proportion <1s** | **N Sequences** |
| Timing of sequence start | 0.80 | 2.29 | 0.88 | 203 |
| Timing of sequence end | 0.82 | 1.96 | 0.79 | 203 |

To obtain additional information on social attention, sequences with other capuchins present besides the tool user were coded again by one coder (ZG) following a different protocol (Table S3). Per sequence, we coded the age-sex class of each other visible capuchin, as well as the age-sex class of capuchins paying social attention to, scrounging from, or displacing the tool user. We identified capuchins whenever possible. Juveniles were usually not able to be sexed, as to do so requires close up images of their genitals for which the quality of the camera traps and image angle are insufficient. We also did not observe any subadult females present during tool use sequences, so only coded subadult males.

### Statistical analyses

All statistical analyses were done in R v. 4.3.1 (Team, 2022). All Bayesian regression models were fit via the “brm” function in the brms package v. 2.16.1 (Bürkner, 2017). We calculated contrasts to evaluate the credibility of the difference between categories (e.g., comparing juveniles’ time to open an item to subadults’ time) using the “hypothesis” function in brms. To identify credible effects, we examined the proportion of the posterior probability (PP) for each contrast that exceeded 0. We considered effects with PP *>*80% as moderately reliable and PP *>*95% as strongly reliable. Conversely, effects with PP *<*20% or *<*5% indicate similarly reliable effects in the opposite direction. In the aforementioned example, a posterior probability of 98% would reflect that the model estimates a 98% probability of juveniles taking more time than subadults to open an item.

#### Proficiency

To compare tool use proficiency between age classes, we first subset our data to only sequences where the item was successfully opened, as failure was rare, occurring in fewer than 4% of observed tool use events. We further limited our analyses on proficiency to sequences where the tool user could clearly be identified. We also excluded sequences that were split across multiple videos (*<*7 % of all sequences), as several seconds of behaviour could have been missed in between triggers. Additionally, we only included sequences where sea almonds were the item being processed, since they constituted the vast majority (98%) of all tool use sequences and other items were under-represented. In our dataset, each row represented a tool use sequence and variables were aggregated to the sequence level.

We quantified tool-using proficiency per sequence in four different ways: i) seconds needed to open an item, ii) number of pounds to open an item, iii) number of repositions of the item on the anvil in between strikes and peels of the sea almond’s exocarp, iv) number of misstrikes and strikes that sent the item flying off the anvil. To compare sequence duration between age classes, we fit a generalized linear Gamma regression (*Model e1*), with sequence duration in seconds as the outcome. As predictors, we included the age class of the tool user (juvenile, subadult, or adult, with juvenile as reference level), the colour of the sea almond (brown, green, red, or unknown, corresponding to its ripeness and with brown as reference level), the material of the anvil (wood or stone, with stone as reference), and a random effect of the identity of the tool user. We included the colour of sea almond and the material of the anvil, as we expected both to potentially have an effect on the number of pounds required to open a sea almond. Fresh sea almond fruits are green, and turn yellow, then red, as they ripen (Perez & Condit, n.d.). After drying, cached fruits turn brown, and we expect these dried fruits to be easier to crack than fresh, green ones. Stone and wood have different densities and hardnesses, likely also affecting how easily the sea almond cracks when smashed with a stone hammer.

To compare number of pounds to open an item between age classes, we fit another generalized linear model (*Model e2*), with the same structure as the model described above. As the predictor was the discreet number of pounds, we used a Poisson distribution. We also compared the rate of pounding (i.e., number of pounds per second) between age classes by running a model with the same structure as *Model e2*, but with an offset of the log of sequence duration in seconds (*Model e2b*). We used the same model structure—without an offset—to consider the number of times an item was repositioned (*Model e3a* and item peels (*Model e3b*) during a sequence. Misstrikes, where a capuchin attempts to strike the item but misses it entirely, were not very common. Thus, to compare its occurrence between age classes, we fit a zero-inflated Poisson model, with the same predictors as above (*Model e4a*). Lastly, to compare the occurrence of items flying off the anvil between age classes, which was similarly rare, we fit another zero-inflated Poisson model (*Model e4b*).

#### Development

To examine the development of tool use proficiency over time, we looked at known individuals and how their processing of one colour of sea almond (brown, which was most frequently processed) changed over the course of the full year of coding. We visually considered how the number of pounds, number of repositions, and number of misstrikes changed, with the expectation that (sub)adults would remain relatively stable, while juveniles would show fewer pounds, repositions, and mistakes over time, as their proficiency increased. We modelled the number of pounds using a Poisson GLM (*model dev1*), with predictors including the interaction between standardized days since the start of the study period and tool user identity. Furthermore, we also considered a case study of one adult male, estimated to be the oldest in the capuchin group, to see whether and how his tool-using behaviour changed since the beginning of sampling in this project (in 2017).

#### Social attention

To examine social learning’s role in tool use acquisition, we ran further analyses on a subset of sequences where other capuchins were present. Here, we also excluded sequences split across several videos, as well as sequences where one of the capuchins present was unable to be assigned an age-sex class, or where the ending of the tool use sequence was not captured on video (so the sequence likely continued after the video stopped recording). We also restructured our dataset to have each row reflect an individual present during a tool-using sequence, with a variable indicating whether they paid social attention yes/no.

First, to consider which age class was more likely to receive social attention, and which age-sex class was more likely to pay social attention to tool use, we fit a generalized Bernoulli logistic regression (*Model socatt1*). The outcome of this model was social attention (yes/no), while the predictors included the age of the tool user (juvenile, subadult, adult, with juvenile as reference) and the age-sex class of the observer (juvenile, subadult male, adult female, and adult male, with juvenile as reference). We also included location (CEBUS-02 or EXP-ANV-01) and two social factors which may affect the probability of social attention occurring, namely the total number of capuchins present during the sequence, and the number of capuchins scrounging during the sequence. Lastly, we included an offset of the log of sequence duration, to account for longer sequences having more opportunities for social attention, and a random effect of sequence ID, to account for non-independence of observations from the same sequence.

Second, to consider whether more efficient tool users, or specific individuals, received more social attention, we subset the data further to only sequences where the sea almond was opened, and where the tool user could be identified. We ran another generalized Bernoulli logistic regression (*Model socatt1b*) with the outcome of social attention yes/no, and predictors of tool user age, age-sex of observer, the number of pounds used to open the item, and the number of mistakes during the sequence. We also included an offset of sequence duration, and also the identity of the tool user as a random effect.

All models were fit with regularizing Normal (0,1) priors for intercepts and predictors, and exponential (1) priors for standard deviations of varying effects. For each model, we did a prior predictive simulation to compare our chosen priors to default priors, and to evaluate the identifiability of parameters. The final models were run with three chains of 3000 iterations each, with a 1500 iterations warm-up per chain. Our models were stable with large effective sample sizes (Bulk ESS and Tail ESS over 1000 for all estimates; (Bürkner, 2017) and Rhat values *<*1.01 (Vehtari et al., 2021)). Additionally, Pareto k estimates for all models were below 0.7. We visually assessed model fit and confirmed our choice of priors using the posterior predictive check function.

## Data availability

Details of model output are available as supplemental information. All code and data necessary to replicate analyses are publicly available on (Goldsborough, 2025).

## Ethical note

We obtained permission for this study from the relevant authority Ministerio de Ambiente, Panama (scientific permit no. SE/A-37-17, SC/A-23-17, SE/A-98-19, SE/A-6-2020, ARB-158-2022, and corresponding renewals and addenda). This project was registered under STRI IACUC SI-25021.

## Results

Between January 2022 and January 2023, we recorded a total of 3644 tool use sequences at the two anvil sites spread across 215 days. CEBUS-02 was utilized more, with 2568 observed sequences compared to 1076 at EXP-ANV-01. Both locations have gaps in tool use activity over the time period (Figure S1). At CEBUS-02, the video cameras ran out of battery prematurely due to the higher activity (cameras ran 2022-01-10 until 2022-06-29 and 2022-07-16 until 2022-09-30). At EXP-ANV-01, the capuchins only regularly started using the novel anvil in March 2022, and reduced their activity after a tree branch fell on the anvil on 2022-10-10, though cameras ran the full time period (2022-01-10 until 2022-07-16 and 2022-07-17 until 2023-01-01). The average number of sequences observed on a day when tool use occurred was 16.95 (range 1-114). Sea almonds were the item being processed in the vast majority of tool use sequences (3578 sequences equal to 98%, Table 3).

**Table 3:**
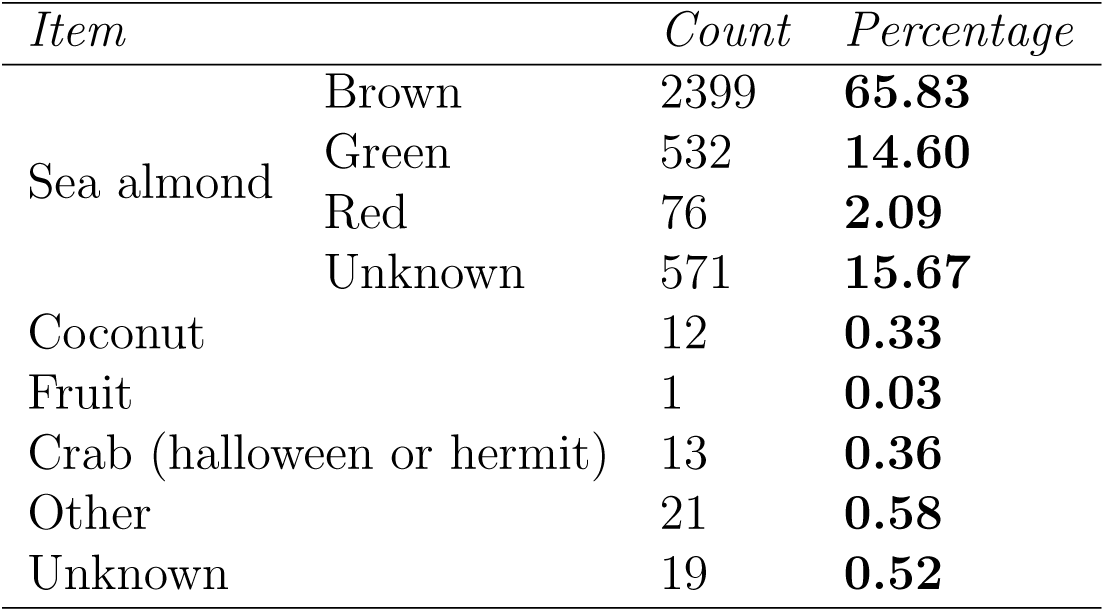
Frequency of items being processed during all observed tool use sequences.

Capuchins successfully opened the item in 89% of the sequences. Success rates were lower for juveniles (79%) compared to subadults and adults (93% and 91% respectively). The number of pounds needed to open a single item ranged from 1 to 34. In the remaining sequences, the tool user either abandoned the item (4%), relocated to another anvil with the item (3%), or the ending was not captured on camera (4%). At each sites, a single hammerstone was used in over 90% of the tool use sequences, and all hammerstones were used by all age classes.

We were able to classify the tool user’s age class in all of the sequences, and reliably identify the tool user in 98% of the sequences (see Table 1 for the number of tool use sequences observed per individual). The tool user was the only individual visible in the video in 70% of all sequences. When other capuchins were present (1107 sequences), we observed social attention occurring in 222 sequences (20%). Both displacements and scrounging were rare, occurring respectively in 12% and 22% of sequences in which other capuchins were present.

### Tool-using proficiency

#### Efficiency (number of pounds and processing time)

The sample used for analyses consisted of 2879 sequences where: i) the tool user successfully opened a sea almond, ii) the tool user was identified, and iii) the sequence was completed in a single video (n_adult_ = 663, n_subadult_ = 1606, n_juvenile_ = 610). Our models estimated that juveniles need considerably longer to open a sea almond with stone tools than subadults and adults (Figure 2, for model estimates see Tables S4 & S5). Juveniles required an average of 19.49 seconds (95% CI [14.44, 26.05]) and 5.42 pounds (95% CI [4.48, 6.36]) to open an item, considerably longer than subadults, who took on average 9.78 seconds (95% CI [4.48, 20.49]) and 3.78 pounds (95% CI [2.34, 5.81]), and adults, who took 11.25 seconds (95% CI [5.16, 24.53]) and 3.94 pounds (95% CI [2.44, 6.05]). Juveniles taking more seconds and more pounds to open an item than subadults and adults were credible effects (all PP *>*0 = *>*98%). Subadults and adults did not differ much from one another in either model.

**Figure 2:**
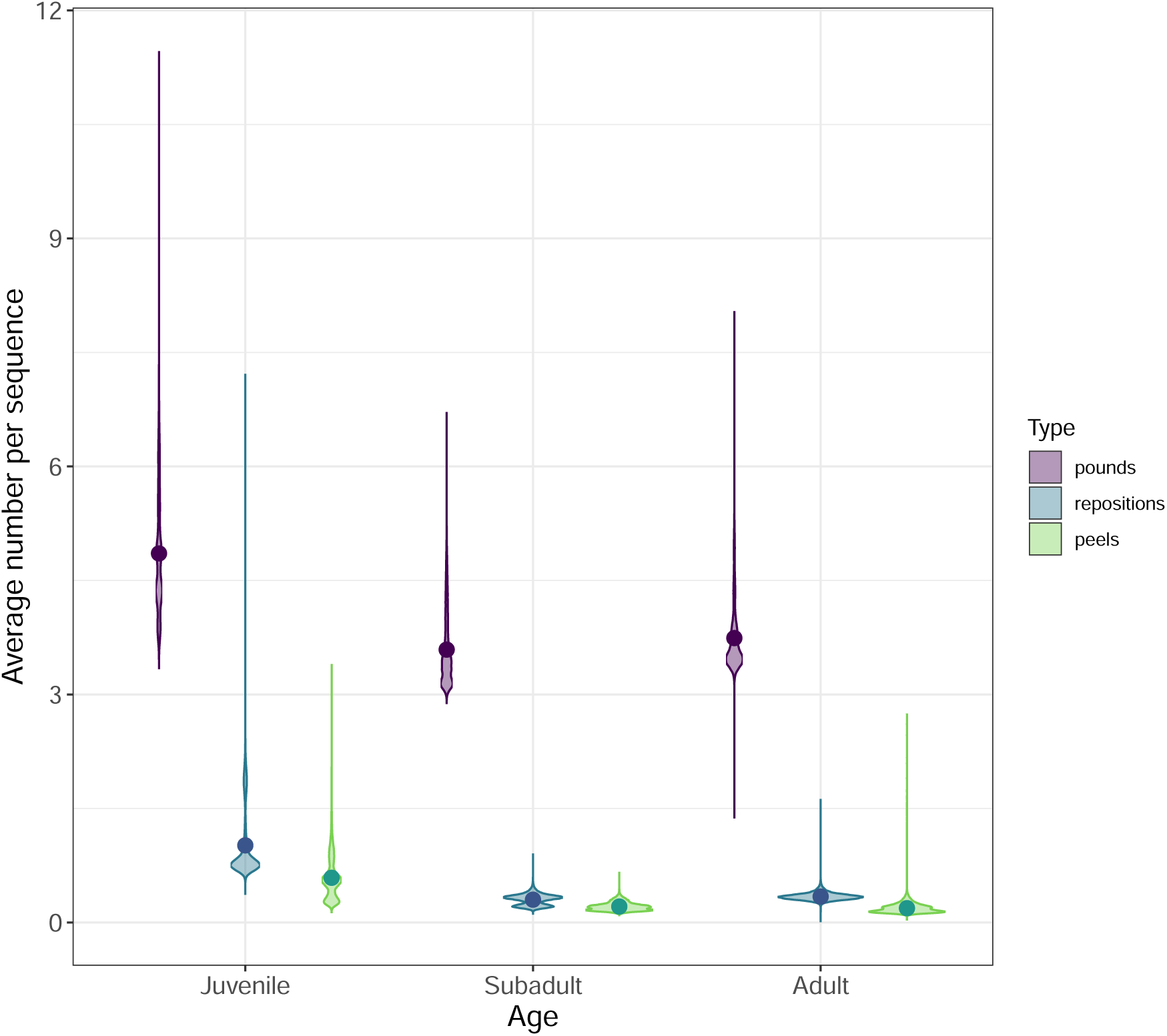
Model estimates and mean values of pounds, repositions, and peels in one tool use sequence opening a single sea almond per age-class (juvenile, subadult, adult). Model estimates are reflected by violin plots, and observed means from data are represented as points.

The level of ripeness (colour) of the sea almond and anvil material affected both seconds and pounds required to process a sea almond. Fresher sea almonds (green and red in colour) took more time and more pounds to open than older (brown) sea almonds. We estimated green almonds to require 1.32 times more seconds (95% CI [1.27, 1.39], PP *>*0 ≈ 100%) and 1.27 times more pounds (95% CI [1.22, 1.34], PP *>*0 ≈ 100%) than brown almonds. Red almonds required 1.49 times more seconds (95% CI [1.34, 1.67], PP *>*0 ≈ 100%) and 1.32 times more pounds (95% CI [1.21, 1.46], PP *>*0 ≈ 100%) than brown almonds. Red and green sea almonds did not differ considerably from one another in processing times or number of pounds. The material of the anvil did not affect processing times, but the number of pounds was slightly higher (1.12 times, 95% CI [1.08, 1.16], PP *>*0 ≈ 100%) at the stone anvil than at the wooden anvil. The relationship between the number of pounds and the sequence duration (i.e. the speed at which individuals pound) was positive between age classes (Figure 3), and older individuals pounded faster than younger ones. Juveniles pounded 0.75 times slower than adults (95% CI [0.62, 0.93], PP *>*0 = 98%) and 0.74 times slower than subadults (95% CI [0.61, 0.89], PP *>*0 = 99%, for all estimates see Table S6).

**Figure 3:**
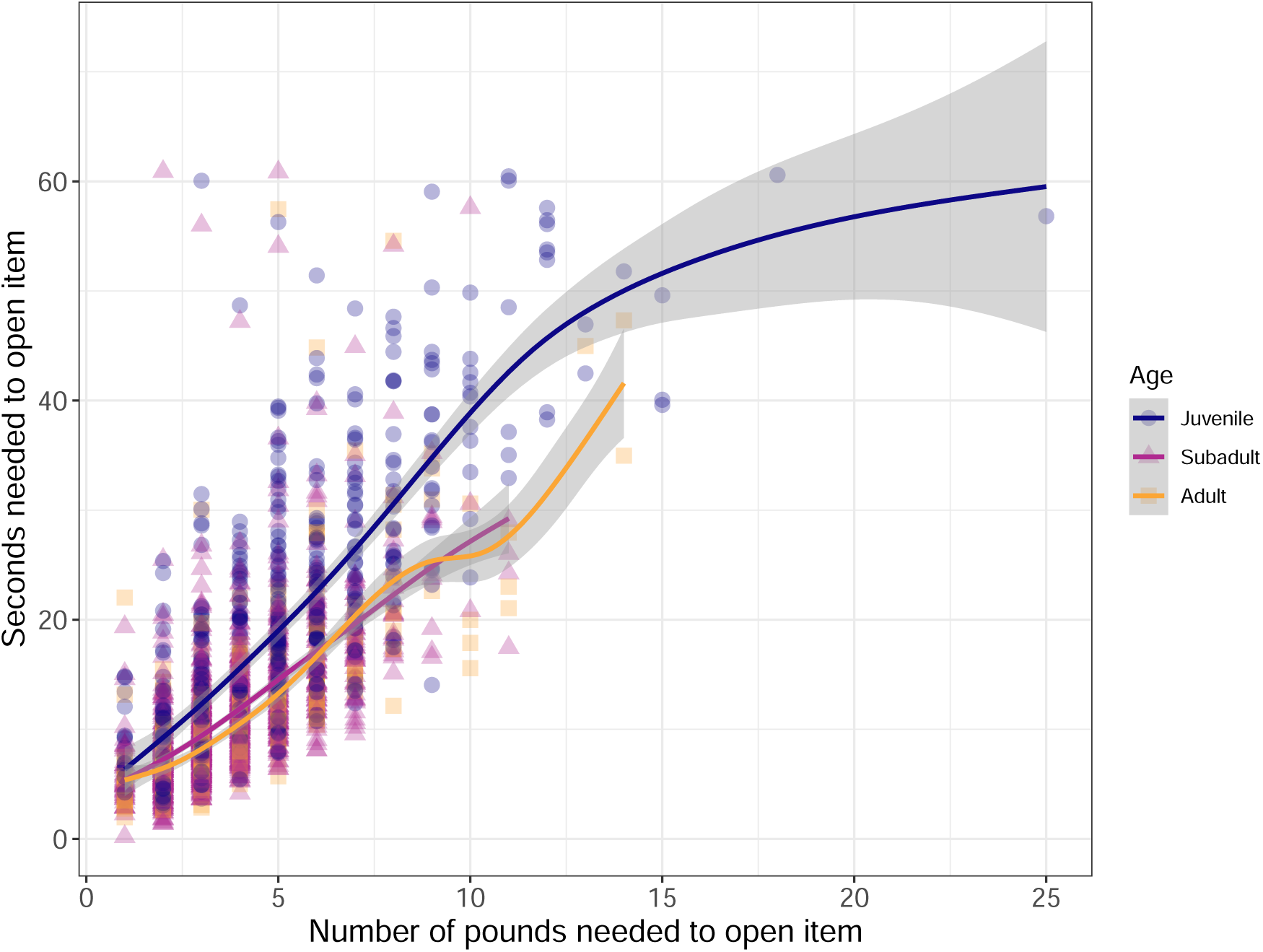
Relationship between number of seconds and number of pounds needed to open a single item using stone tools, separately for each age class.

#### Repositioning and peeling

All age classes showed repositioning of the item on the anvil during a tool use sequence, or picking it up and peeling it before continuing to strike, but juveniles repositioned and peeled more than adults and subadults (Figure 2, for model estimates see Tables S7 & S8). Juveniles showed considerably higher incidences of repositioning and peeling (mean_reposition_ = 1.07, 95% CI [0.62, 1.70]; mean_peel_ = 0.46, 95% CI [0.27, 0.73]) than subadults (mean_reposition_ = 0.32, 95% CI [0.09, 1.23]; mean_peel_ = 0.19, 95% CI [0.06, 0.64]) and adults (mean_reposition_ = 0.33, 95% CI [0.09, 1.21]; mean_peel_ = 0.19, 95% CI [0.05, 0.72]). Juvenile tool users repositioning and peeling more than subadults and adults were credible effects (all PP *>*0 = *>*98%). Tool users peeled fresher sea almonds, in particular green ones, more often than brown, older sea almonds (1.36 times more frequently, 95% CI [1.15, 1.62], PP *>*0 ≈ 100%). The material of the anvil made no difference in the number of repositions or peels.

#### Mistakes

Tool users rarely made visible mistakes during tool use sequences (occurring 301 times in the sample of 2879 tool use sequences), and in 48% of cases the tool user making a mistake was juvenile. The most common type of mistake was the item flying off the anvil after being struck (occurring 153 times), followed by misstrikes (99 times), and dropping the hammerstone (occurring 49 times). Juvenile tool users were more likely to make misstrikes or send the item flying (mean_misstrikes_ = 0.06, 95% CI [0.01, 0.29]; mean_itemflies_ = 0.17, 95% CI [0.10, 0.29]) than subadults (mean_misstrikes_ = 0.02, 95% CI [0.00, 0.50]; mean_itemflies_ = 0.04, 95% CI [0.01, 0.15]) and adults (mean_misstrikes_ = 0.03, 95% CI [0.00, 0.83]; mean_itemflies_ = 0.03, 95% CI [0.01, 0.12]). Juveniles making more mistakes than subadults was a credible effect (misstrikes PP *>*0 = 91%, items flying PP *>*0 = 99%), as was juveniles sending more items flying off the anvil than adults (misstrikes PP *>*0 = 75%, items flying PP *>*0 = 100%; for model estimates see Tables S9 & S10).

The material of the anvil also affected the likelihood of the item flying off: a sea almond was 3.35 times more likely to fly off of a stone than wooden anvil (95% CI [2.46, 4.57], PP *>*0 = 100%). Green and red sea almonds were also 3.63 (95% CI [2.61, 5.05], PP *>*0 = 100%) and 2.32 times (95% CI [1.15, 4.48], PP *>*0 = 98%) more likely to fly off when struck than brown, dried out, sea almonds.

### Tool use development

We observed little change in tool use proficiency over the year in any age class. Adults’ performance was generally stable over time, and juveniles and subadults likewise showed minimal changes in the number of pounds. Only one individual, TER, showed a modest improvement over the study period (estimated slope = –0.08, 95% CI [-0.19, 0.05].

Notably, he had been the least proficient juvenile at the start (for model estimates, see Table S11; individual trajectories shown in Figure S2. Adults’ performance was the most stable, but juveniles and subadults also did not show considerable changes in the number of pounds (for model estimates see Table S11), repositions, and misstrikes over time (Figure 4). The reduced sampling near the end of the observation period, combined with some individuals (mostly adults) showing clustered activity rather than activity spread over time, means that there was more uncertainty in our estimates at particular times of the year.

**Figure 4:**
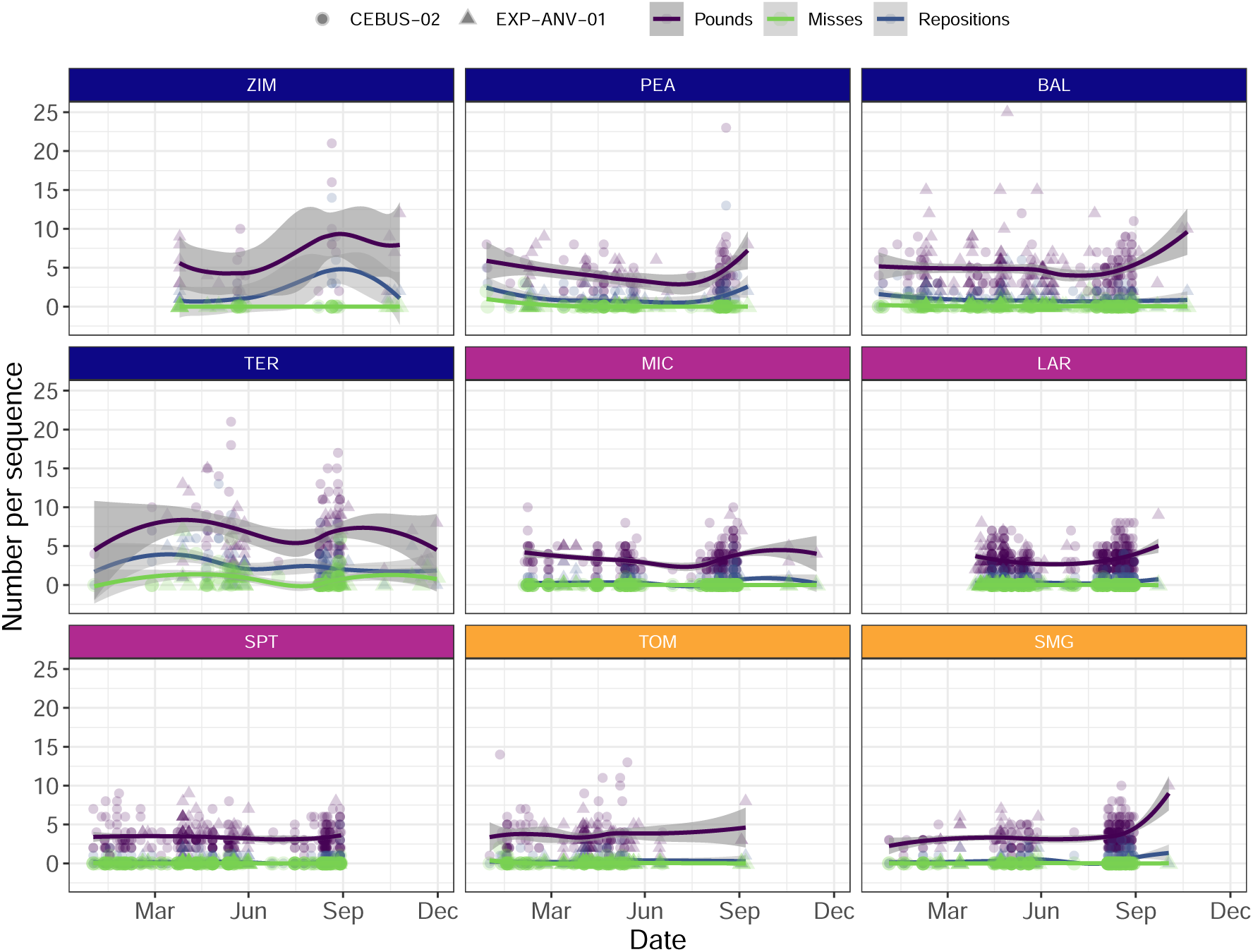
Development of number of pounds (purple), misstrikes (green), and repositions (blue) per individual over the full observation period for brown sea almonds only. The colour of the box with the individuals code represent their age class (blue for juveniles, lilac for subadults, orange for adults). Points are raw data, with point shape corresponding to anvil location. The smooth curve is estimated using method “loess” and formula “y_x”.

Importantly, one group member—ABE—who was present in the group since the beginning of the project, was never observed to use tools in the 1-year period comprising this study. This adult male is estimated to be at least 30 years old, and the likely alpha male of the group, inferred by his size, his recruitment by other group members in conflicts, and the fact that we never observed him receiving aggression from any other group members. We observed him using tools 73 times in data collected 2017-2019, yet we never saw him use tools in the current observations. However, ABE displaced a tool user from the anvil 14 times (out of 176 total displacements), and then consumed the item they opened. No other individuals aggressively scrounged from tool users in this manner.

#### Social attention

After filtering, our sample contained 907 sequences that were completely captured within a single video, during which other capuchins aside from the tool user were present, who could all be assigned an age-sex class. The majority of other capuchins present were juveniles (1034/1652 capuchins present, 63%), followed by subadults (326, 20%) and adult males (165, 10%). Adult females were rarely present during tool use sequences (127 times, 8%). In this filtered sample, social attention occurred in 189 (21%) of the sequences.

Individuals paying social attention were overwhelmingly juveniles. An adult male was observed paying social attention once, adult females five times and subadult males 20 times. Of the 215 occurrences of juveniles paying social attention, 23 were identified as known tool-using juveniles. The remaining unidentified juveniles were likely not tool users, as many were too young (*<*3 years) to easily be identified and we have not observed regular tool use by any individuals of this age.

Our model exploring factors predicting the probability of social attention also found juveniles to be the age class most likely to pay social attention. Given an adult male tool user at the most frequently used location (CEBUS-02), juveniles had a probability of showing social attention of 0.14 (95% CI [0.08, 0.22]), considerably higher than subadult males (0.04, 95% CI [0.02, 0.09]), adult males (0.02, 95% CI [0.01, 0.04]), and adult females (0.02, 95% CI [0.01, 0.06]). In terms of who received social attention, given a juvenile observer at CEBUS-02, juveniles were less likely to receive social attention [0.07, 95% CI: 0.04-0.10], than subadults [0.17, 95% CI: 0.13-0.22] and adults [0.14, 95% CI: 0.08-0.22], who did not differ significantly from one another (Figure 5). Juveniles being more likely to give and less likely to receive social attention than other age-sex classes was a credible effect (PP *>*0 for all contrasts = 100%).

**Figure 5:**
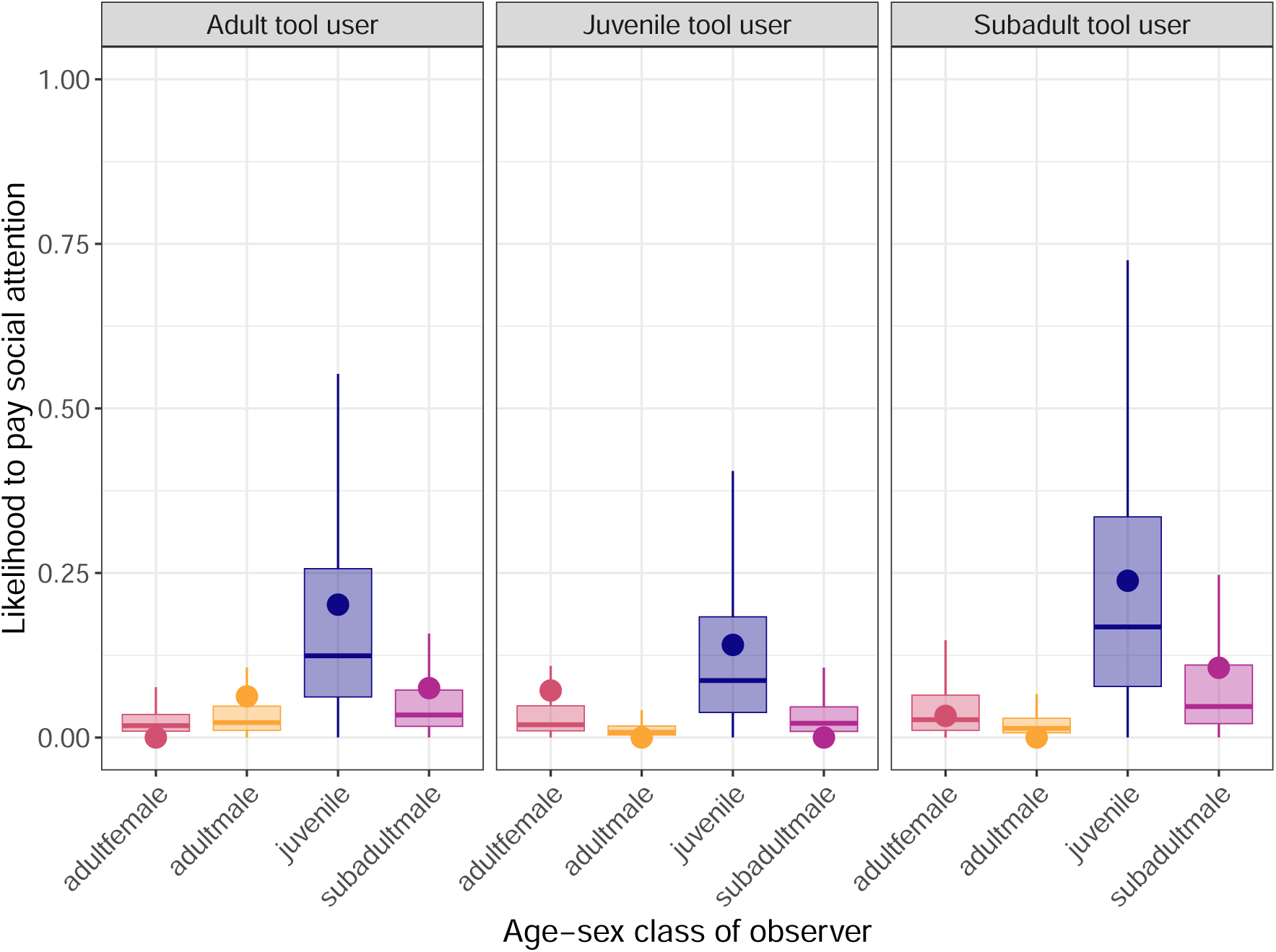
Likelihood of paying social attention to a tool user, presented per age-sex class of the observer and age of the tool user. Model estimates are reflected by box plots, and observed means from data are represented as points.

When more capuchins were present during a tool-using sequence, the chance of social attention occurring was lower. However, in sequences where more individuals were scrounging, social attention was more likely to occur (Figure 6, for model estimates Table S12). When we considered the type of scrounging (whether it was tolerated by the tool user during the tool use event, or only occurred after the tool user had left), social attention mostly occurred in combination with tolerated scrounging (tolerated scrounging occurred in 111/209 sequences with social attention, compared to scrounging afterwards occurring in 14/209 sequences). Although sequences with other capuchins present were rarer at EXP-ANV-01 (180) than CEBUS-02 (664), social attention was more likely at EXP-ANV-01 (Table S12). Given the most frequently occurring combination of a juvenile paying attention to a subadult tool user, the predicted likelihood of social attention at EXP-ANV-01 was 0.33 (95% CI [0.24-0.42] vs. 0.17 at CEBUS-02 (95% CI [0.13, 0.22]). Paying social attention was not entirely without risks; in one tool use sequence a juvenile observer got too close to the tool user, and was hit on the foot with the hammerstone (Video S1).

**Figure 6:**
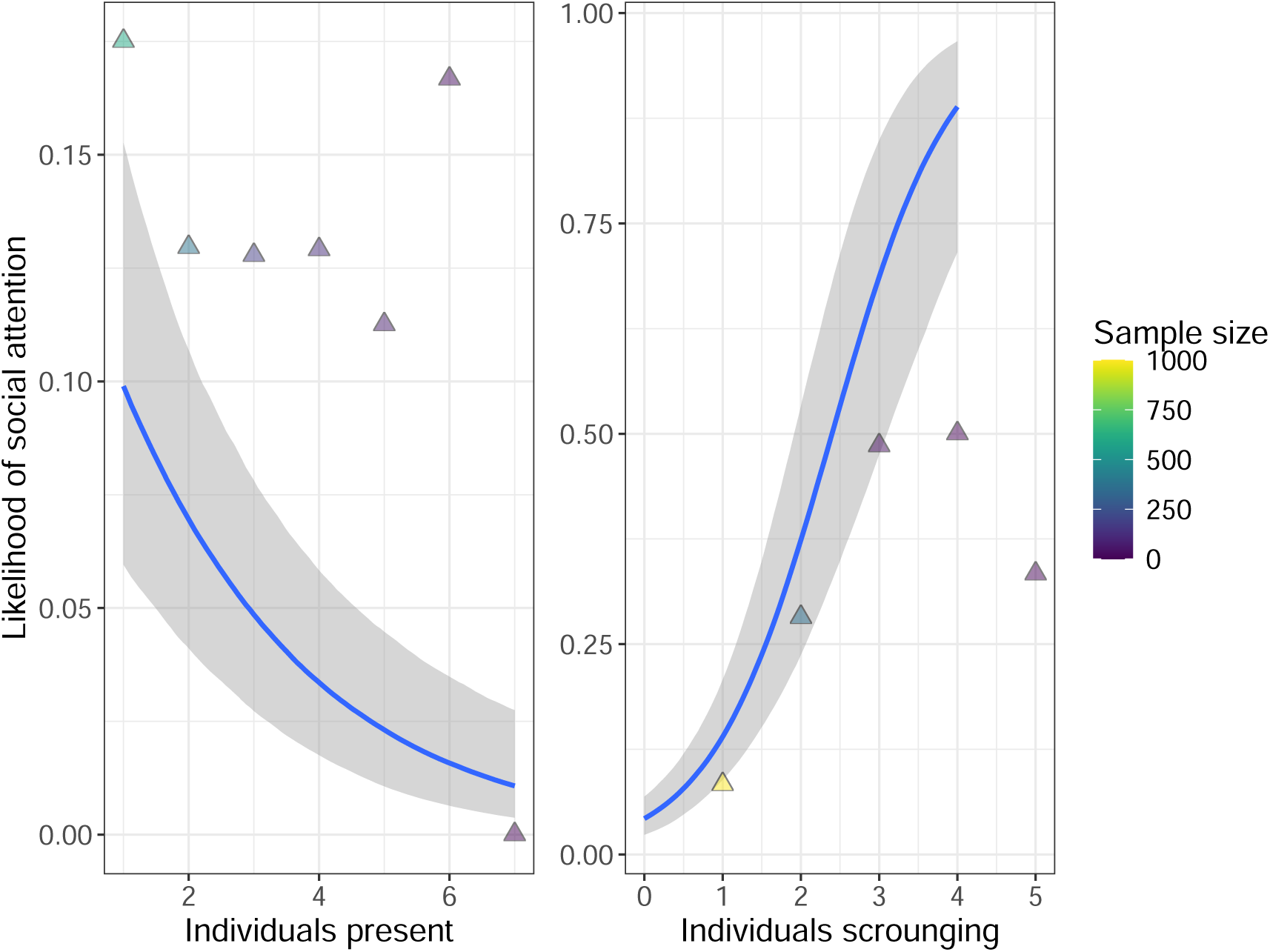
Likelihood of paying social attention to a tool user, depending on number of capuchins present (left) and scrounging (right) in the sequence. Model estimates are reflected by the lines, and observed means from data are represented as points. colour of points reflects the size of the sample contributing to the mean.

To consider whether social attention is preferentially paid to more proficient tool users, we limited our sample to sequences where: a) the item was successfully opened, b) we could confidently identify the tool user, and c) the tool user had more than 10 tool-using sequences in the sample (n = 1240 sequences). Our model estimated a slight trend of more social attention occurring in more efficient tool-using sequences (where fewer pounds needed to open the item), and this relationship was strongest if observers were juveniles (see Figure S3 and model estimates Table S13). Given a juvenile observing a subadult tool user, the likelihood of the juvenile paying social attention was predicted to decrease from 0.31 (95% CI [0.24, 0.41]) at one pound to 0.14 (95% CI [0.04, 0.35]) at 18 pounds.

The individuals who received most social attention during tool use were the three subadult males, who are all proficient tool users and also showed highest incidence of tool use with other individuals present (Figure 7). Individual subadults differed little from each other in how much social attention they received.

**Figure 7:**
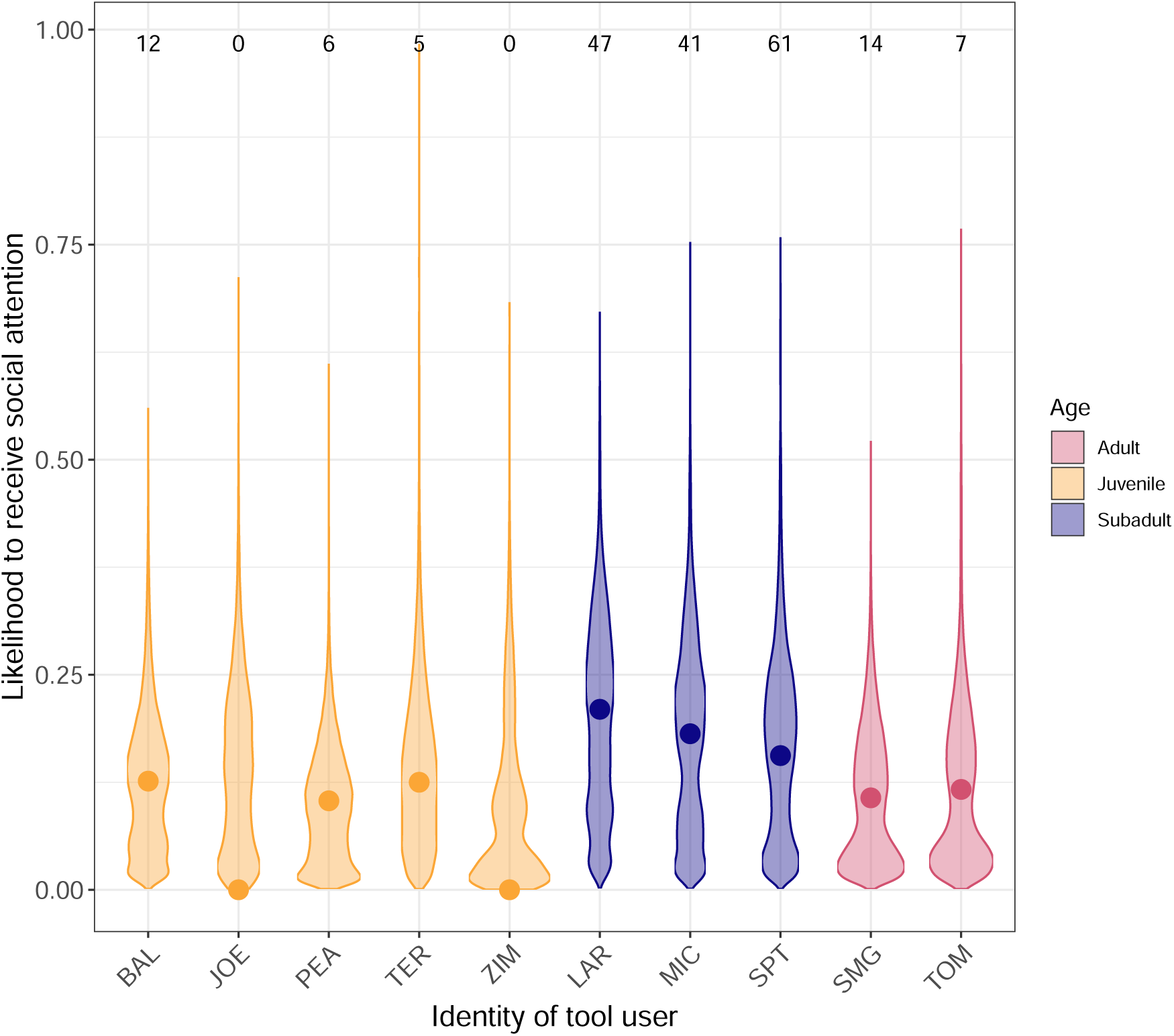
Likelihood of receiving social attention for different tool users. Model estimates are reflected by violin plots, and observed means from data are represented as points. colour reflects the age class of the tool user, and the numbers above the number of sequences with social attention (top) compared to the number of sequences with other capuchins present where no social attention occurred (bottom in italics).

## Discussion

Stone tool use by white-faced capuchins on Jicarón is a complex behaviour for which proficiency takes years to develop. Compared to subadult and adult tool users, juvenile tool users need more time and more pounds to open an item using tools. Juveniles more frequently reposition the item in between strikes, and make more mistakes. Subadults and adults show comparable levels of proficiency on all measures. Four identifiable tool-using juveniles did not markedly improve in tool use proficiency over the course of approximately one year. No other capuchins were present in the vast majority of tool use sequences and social attention to the tool user was rare (occurring in just 6% of all sequences). Nonetheless, we uncovered some robust patterns in who paid social attention to whom, and in what context. Juveniles were most likely to pay social attention to tool use, and most of these juveniles were too young to be tool users themselves. Social attention was directed at subadults and adults, with a slight preference towards more efficient tool users. Subadults received the most social attention. The more capuchins were present during a tool-using sequence, the lower the probability that social attention occurred. Scrounging showed the opposite relationship: the more individuals scrounging, the higher the chance of social attention.

### Differences in tool use proficiency

It is unsurprising that juveniles are less proficient tool users than adults and subadults, regardless of the measure of proficiency considered. This pattern is similar to what has previously been described in macaque stone tool use (Tan, 2017) and for this species of capuchin in other extractive foraging tasks (Barrett et al., 2017). However, juveniles were still very successful in opening items, and the difference in their efficiency compared with subadults and adults is small. This highlights an important distinction between basic success (being able to open a nut most of the time) and the maximum level of proficiency observed in adults, which likely reflects a combination of physical maturation and skill refinement. Overall, white-faced capuchins had a slightly higher success rate than what is reported in studies of other nut-cracking primates. The item being abandoned entirely only occurred in 4% of all sequences, much lower than in chimpanzees (failure occurring in 8% of tool use events (Berdugo et al., 2024)) and robust capuchins (failure in 2-38% of events depending on population (Falótico & Ottoni, 2016; Falótico et al., 2024; Spagnoletti et al., 2011)). This does not necessarily reflect higher skill of the white-faced capuchins, but could also be an artifact of their high persistence, motivation and tenacity. It is important to note that we excluded tool use sequences that spanned more than one camera trap video from analyses. The age distribution in these excluded sequences was comparable to that of the sequences retained for analysis. The exclusion of split sequences is unlikely to bias our results, because sequence length and processing speed are age-dependent: juveniles are more likely to be less proficient and have not completed an item by the end of a video, while (sub)adults often process more quickly and may start a second (or third) item at this time.

The clearest distinction between juvenile and (sub)adult tool use proficiency was the much higher incidence of mistakes by juveniles. Mistakes were rare in our sample overall, similar to bearded capuchins (Fragaszy et al., 2020), but, contrary to the findings on bearded capuchins, we found that juveniles were considerably more likely to miss the item, or have the item fly off the anvil. Juveniles also more frequently repositioned the food item between strikes. More frequent repositioning by juveniles could reflect poorer control over the hammerstones than adults. This ties in with our surprising finding that at both sites, a single hammerstone was used in over 90% of the sequences, despite multiple other hammerstones being provided. These preferred stones were large and heavy, and thus easier to handle for (sub)adults than juveniles. The (lack of) hammer selectivity relative to body size at the experimental anvils is an interesting avenue for future research, as it stands in stark contrast to findings in chimpanzees (C. Boesch & Boesch, 1983), macaques (Gumert & Malaivijitnond, 2013), and robust capuchins (Visalberghi et al., 2009). The preference for larger, heavier hammerstones for use in nutcracking appears to be widespread in this population (Carlson et al., in prep). This may be related to the fact that for certain tasks, large hammers might yield large dividends for gracile capuchins, which are the smallest, lightest tool-using primates.

The same hammerstone may have been used so consistently at each site because there was also little variation in the food items being opened. In nearly all sequences, capuchins processed sea almonds. However, the level of ripeness of the sea almond did appear to affect ease of processing: green, fresher, sea almonds required more seconds and more pounds to open than brown sea almonds, were peeled more frequently, and were more likely to fly off the anvil when struck. This likely reflects the harder exocarp of the fruit at this stage, before it is dries out and becomes more brittle. The lack of hammerstone selectivity for sea almond ripeness and higher incidence of peeling with green sea almonds indicates that perhaps the additional step of peeling off the exocarp in between strikes helps overcome the difference in hardness, a technique which juveniles might also need to learn to become a more proficient tool user. However, juveniles also peeled more frequently than (sub)adults, so it is also possible that juveniles lack the knowledge/ability to precisely strike the green sea almonds without them flying off the anvil – which green sea almonds are more likely to do — and that peeling the fruit makes it easier to strike. In either case, becoming a proficient stone tool user might not only involve learning *how* to process, but also *what* and *when*. Sea almonds have variable fruiting seasons, and although we know little of seasonal fluctuations in food availability on Jicarón, our data do show sea almond processing peaking bi-modally (Figure S1). Additionally, (sub)adults seem to focus more of their tool-using efforts at specific times of the year (see Figure 4), whereas juveniles use tools at a more consistent rate throughout the year. Future studies can elucidate the cause of these seasonal fluctuations in tool use, and also explore whether adults indeed concentrate their tool-using efforts when sea almonds are most abundant or, alternately, when other, easier to consume food resources are unavailable.

### Development of tool proficiency

The lack of change in tool use proficiency by juveniles over the course of our study period suggests that honing these skills to adult levels of proficiency is likely a slow process, or one limited by physical features (like strength) which only mature in adulthood. White-faced capuchins are a long-lived species, with a lifespan up to 50 years (Fragaszy et al., 2004), similar to chimpanzees, where it has been shown that becoming a proficient nut-cracker can take 3-7 years of practice (C. Boesch & Boesch, 1990; Matsuzawa, 1994). Our results, though not yet sufficiently longitudinal for a full understanding of tool use ontogeny, suggests that in white-faced capuchins, becoming a proficient stone tool user may be a similarly extended process. Furthermore, our finding that the oldest tool-using male in the group, ABE, did not use tools at all during the 1 year period of this study, hints at a potentially upper age limit for stone tool use in white-faced capuchins, as has been documented in chimpanzees (Howard-Spink et al., 2024). ABE is the only individual to displace tool users from the anvil to steal the item they opened, which suggests he is still interested in the resources, but perhaps no longer able or willing to invest the energy and time to use tools himself. Here, too, a more longitudinal approach would be very valuable to examine the fluctuations in tool use proficiency across the lifespan.

### The role of social attention in tool use acquisition

Stone tool use at the experimental anvils appeared to be a largely solitary activity. As such, opportunities for social attention seem limited, and even when other capuchins were present, social attention only occurred in 20% of the sequences. Data from camera traps may underestimate social attention, as individuals out of frame could be looking at the tool user without this being captured. However, attention from a distance likely only provides the observer with coarse information on the tool use behaviour. The low frequency of close social attention in our sample is surprising compared to other nut-cracking primates. In one study on tufted capuchins, conspecifics observed 36% of all nut-cracking events (Coelho et al., 2015), and in chimpanzees, juveniles observe hundreds of nut-cracking events from their mothers (Estienne et al., 2019). Yet on Jicarón, juvenile capuchins have no opportunity to learn tool use from their mothers, as females do not use tools (Goldsborough et al., 2024). Individuals paying social attention to tool use were mostly juveniles too young to be tool users themselves, suggesting that social observation may be most important for the acquisition of the tool use behaviour (Barrett et al., 2017). Although scrounging was not as common as in tufted capuchins (Coelho et al., 2015), the more individuals were scrounging, the higher the likelihood of social attention occurring. Furthermore, when scrounging did occur together with social attention, it was nearly always tolerated by the tool user. The tool users who received the most social attention, subadult males, are also known to be tolerant of juveniles (Jack & Riley, 2014). Our results echo the idea that inter-individual tolerance plays a crucial role in social learning of complex behaviours (van Schaik et al., 1999). Furthermore, capuchins might also rely on local enhancement and interaction with temporally enduring artifacts of tool use to learn about the tool use behaviour (Fragaszy et al., 2013). In the absence of many opportunities for direct social learning, debris left on the anvil after a tool use sequence, such as partially processed food items, and wear on the hammerstone and anvil may provide important information to individuals visiting the site. Capuchins on Jicarón using this physical evidence of tool use as a cue could also explain the “burn-in period” at the novel anvil, EXP-ANV-01, since perhaps regular tool use by many individuals only started once enough debris had accumulated from sporadic tool use events, marking this location as a suitable anvil for tool use.

### The importance of location

Here, we report on data collected from only two anvils in the tool-using groups’ range. However, we have identified at least 10 different anvils that are habitually used for stone tool use, and over 300 ephemeral locations. While CEBUS-02 is the most frequently used anvil, which has, to the best of our knowledge, been in use for the longest time (at least since 2004), it is possible that most social learning of tool use occurs elsewhere, for instance during tool use in the intertidal zone (Goldsborough et al., 2023). We do see a difference in the likelihood of social attention between the two experimental anvils: CEBUS-02 has more opportunities for social attention, but social attention is more likely at EXP-ANV-01. This could be due to CEBUS-02 being an established, well-used anvil prior to the current study. We infer that this anvil is used so frequently because it is located in a productive foraging area (next to a stream), leading to more capuchins being present at the same time. In contrast, we purposefully placed the experimental anvil EXP-ANV-01 in a location without evidence of prior tool use, meaning this location likely held less interest for the capuchins. This inference is supported by the fact that it took capuchins taking several months to start regularly using this anvil. The localized availability of ripe sea almonds at this location may have also influenced the latency to capuchins visiting this anvil. Another, non-exclusive, possibility is that, either due to anvil material or other properties of the site, CEBUS-02 is simply a “better” anvil. In this case, there may be more competition over this site, and lower tolerance for observers. We found that on average, capuchins used more pounds at the stone anvil than the wooden anvil, and that items were more likely to fly off the stone than the wooden anvil, further supporting that the wooden anvil at CEBUS-02 has better properties for tool use. Anvil selectivity by tool users and the notion of “wear in” time of new anvils is another interesting avenue for future research, in order to examine what exact physical properties drive these differences in tool use efficiency.

### Spread of tool use to other groups

The seemingly slow mastery of stone tool use by white-faced capuchins also has implications for the spread of this behaviour. Despite being persistent over time, the stone tool use tradition on Jicarón is localized (Barrett et al., 2018). In mainland populations of white-faced capuchins, males are the dispersing sex and first dispersal occurs on average at 4.5 years of age in Santa Rosa (Jack et al., 2012) and 7 years of age in Lomas Barbudal (Perry et al., 2012). On Jicarón, males of this age would not yet be proficient tool users based on our data, and as such, might cease using tools post-dispersal, limiting the spread of the behaviour. Moreover, females on Jicarón do not use stone tools despite being physically capable of it (Goldsborough et al., 2024), this sex bias further constrains potential transmission between groups. However, it is unclear whether the typical pattern of male dispersal holds on Jicarón, or if, instead, both sexes, or only females disperse. Switches in dispersal tendency are known to occur under high density conditions (Clutton-Brock & Lukas, 2012). It appears likely that at least some males remain in the natal group, as we have tracked some individuals from juvenile to (sub)adulthood. Further elucidating the dispersal tendency of these capuchins is crucial to better understand the maintenance and spread of the tool use behaviour.

## Conclusion

We present a first description of variation in and development of stone tool use proficiency in white-faced capuchin monkeys. Using camera traps placed at two experimental tool use anvils on Jicarón island, Panama, we collected nearly a year of tool-using sequences, mostly of capuchins processing sea almonds. Similar to other primates that show percussive stone tool use, white-faced capuchin juveniles are less proficient tool users than adults (in terms of efficiency and technique), and development of proficiency appears to be a slow process. Ripeness of the sea almond affected processing times and efficiency, with less ripe green sea almonds requiring more time and peeling of the skin in between pounds, and generating more mistakes. In contrast to findings from other tool-using primates, tool-using appeared to be a largely solitary activity, with limited opportunities for social learning and even fewer actual occurrences of close social attention. However, when social attention to tool use did occur, there were clear patterns to it that echoed studies in other primates. Social attention was mostly paid by juveniles to proficient subadults, who were tolerant of scrounging. We also found an effect of location on the likelihood of social attention occurring, suggesting social learning opportunities differ between sites. In short, we lay the groundwork for both comparisons between stone tool-using primate species, as well as future research on other aspects of white-faced capuchin stone tool use like seasonality and hammerstone and anvil selectivity.

## Supporting Information

**Video ethogram**: https://keeper.mpdl.mpg.de/d/0c1b9853f3f342d8b3da/

**Video S1**: https://youtu.be/tvuClVgwLRQ

**Table S1**: Behavioural ethogram of all behaviours coded for every tool use sequence.

**Table S2**: Overview of sequence-level variables coded for every tool use sequence.

**Table S3**: Behavioural ethogram of behaviours coded per sequence with capuchins present.

**Figure S1**: Number of tool use sequences observed per experimental anvil.

**Table S4**: Posterior mean model estimates of Model e1.

**Table S5**: Posterior mean model estimates of Model e2.

**Table S6:** Posterior mean model estimates of Model e2b.

**Table S7:** Posterior mean model estimates of Model e3a.

**Table S8:** Posterior mean model estimates of Model e3b.

**Table S9:** Posterior mean model estimates of Model e4a.

**Table S10:** Posterior mean model estimates of Model e4b.

**Table S11:** Posterior mean model estimates of Model dev1.

**Table S12:** Posterior mean model estimates of Model socatt1.

**Table S13:** Posterior mean model estimates of Model socatt1b.

**Figure S2**: Conditional effects of time on number of pounds by individual from model dev1.

**Figure S3**: Predicted probabilities from model socatt1b of probabilities of social attention occurring, depending on the number of pounds in the sequence.

## Acknowledgments

We are grateful to Evelyn del Rosario-Vargas, Pedro Luis Castillo-Caballero, Eliecer Vega-Patiño, Juan Rojas Garrido, James Chaves, and Tamara Dogandžić for their assistance with fieldwork, and STRI Panama and the Coiba research station for making this research possible. We further thank Lucia Torrez, Katrin Dieter, and Angie Ruiz for their logistical support. This research was supported by the Alexander von Humboldt Professorship endowed by the Federal Ministry of Education and Research awarded to M.C.C. and the Max Planck Institute of Animal Behavior, as well as a L.S.B Leakey Foundation Grant awarded to M.K.W.C. Lastly, Z.G. and B.J.B. received funding in the form of a grant awarded by the Deutsche Forschungsgemeinschaft (DFG, German Research Foundation) under Germany’s Excellence Strategy – EXC 2117 – 422037984.

