## Supplemental Information for "Development and social dynamics of stone tool use in wild white-faced capuchin monkeys"

### 1 Electronic supplements

**Video ethogram**: https://keeper.mpdl.mpg.de/d/0c1b9853f3f342d8b3da/
**Video S1**: https://youtu.be/tvuClVgwLRQ

### 2 Behavioural ethograms

#### 2.1 Detailed coding of tool use sequences

Table S1: Behavioural ethogram of all behaviours coded for every tool use sequence.

| behaviour | Modifiers | Modifier Options | Definition |
|---|---|---|---|
| pound | | | *The capuchin hits the item with the hammerstone successfully, code pound at (approx.) the moment the hammerstone hits the item.* |
| | pound type | 1. Crouching<br>2. Standing<br>3. Jumping | We differentiate between 3 pound types.<br>1. "crouching" pound, if the legs of the capuchins are at a 90-degree angle.<br>2. "standing" pound, if the legs are extended beyond 90 degrees (and often the body is elongated too).<br>3. "jumping" pound, if both feet leave the ground. |
| | position | 1. 1 foot<br>2. 1 hand<br>3. tail support<br>Multiple selection possible | Modifiers describing the position of the capuchin. The default is 2 footed (both feet on the ground) and 2 handed (both hands on the hammerstone). Only code tail support if you can see the capuchin clearly using their tail (i.e., gripping something with it or putting weight on it). |
| | hammer | 1. Overhead | Modifier to indicate that the hammer was held above the capuchin's head during the hit, so whether any part is higher than their head at pound peak. |
| reposition | | | *When the capuchin repositions the item on the anvil (so anytime they touch/move the item in the sequence) or clearly changes their grip on the hammerstone. Code every time it occurs, but not every movement (e.g. if they grab the item and move it 3 times rapidly, code reposition once).* ***Note:*** *don't code the first time placing/adjusting item or gripping hammerstone at sequence start.* |

| | | | |
|---|---|---|---|
| | object type | 1. hammer<br>2. item<br>3. peel | Whether it is the hammer or item that is repositioned. The third option is that they "peel" the item by holding it in their hand and manipulating it with hands or teeth (often tearing pieces off). **Note**: don't code peel when they do it at the end and eat the item straight after without more pounds. |
| misstrike | | | *When capuchins do not hit the item as intended, or something else goes wrong. You can often use audio cues to hear if they strike the item or the anvil.* ***Note:*** *if capuchin does hit the item successfully but also has a misstrike (e.g. item flies off) then code both the pound and the misstrike.* |
| | mistake type | 1. item flies off<br>2. drop hammerstone<br>3. hammer break<br>4. anvil break<br>5. other | Specify what occurred, either no hit at all (the default), the item flies off the anvil, the hammerstone falls out of their grip, the hammer breaks or the anvil breaks. If not in this list then select other and add a comment to the behaviour. **Important**: also add a comment if you see a capuchin injure themselves or others with the hammer. |
| hammer switch | | | *When a capuchin grabs another hammerstone during the sequence and continues with it.* |
| | hammer location | 1. On anvil<br>2. Off anvil within reach<br>3. Off anvil walk<br>4. Carry in (c) | The location of the hammerstone they switch to. |
| | hammer ID | Enter the hammerstone ID as a comment, or if it's not marked/ unidentifiable:<br>7. Unmarked<br>8. Unknown | Enter the ID of the hammerstone switched to as a comment, which can be one of the marked hammerstones or in the case of an unmarked hammerstone code "unmarked" but describe it. |
| anvil switch | | | *When individuals switch from one anvil to another that are both within view of the camera, within a sequence. So they continue to process the same item at a different anvil.* |
| | anvil material | 1. wood<br>2. stone | The material of the anvil that they switched to. |

Table S2: Overview of sequence-level variables coded for every tool use sequence.

| Variable | Modifiers | Modifier Options | Definition |
|---|---|---|---|
| sequence start | | | *Start of tool use sequence, defined as the moment when the capuchin first places the item on the anvil. If the item is already on the anvil at the start of video, the start of the sequence equals the start of the video.* ***Note****: if capuchins are processing items on another anvil than the experimental anvil (e.g. a branch) make a comment saying "wooden anvil"* |
| | item type | 1. Sea almond green (includes yellow)<br>2. Sea almond brown<br>3. Sea almond red<br>4. Sea almond unknown<br>5. Halloween crab<br>6. Hermit crab<br>7. Coconut<br>8. Fruit (add comment if known)<br>9. Other (add comment)<br>10. Unknown | Code which item is being processed |
| sequence end | | | *The end of the tool use sequence, defined as the moment the capuchin starts consuming the item (if opened) or otherwise when they let go off the hammerstone with no further strikes on the item.* |
| | outcome | 1. Opened<br>2. Relocated<br>3. Abandoned<br>4. Continued | Add modifier to specify how the sequence ended. If the capuchin is eating the item, code it as "opened". The other options are the capuchin taking the item elsewhere ("relocating"), or "abandoning" it. "Continued" is an indicator that the sequence is not yet finished when the video ends. In case the video ends and you don't know what happened to the item (i.e., it's not recorded), keep seqend at "none" and add a comment saying it was unknown/missed. **Note**: if they open the item but do not eat it, then code "abandoned" with a comment saying "opened but not eaten". |
| | scrounging | 1. scrounging<br>2. no scrounging | Code this behaviour if any scrounging (other individuals eating parts of the item opened by the tool user) occurred during or after the sequence. **Important**: only code "no scrounging" if other individuals were present but there was no scrounging. If no other capuchins are visible, then choose "none". |

| | | | |
|---|---|---|---|
| | displacement | 1. No displacement<br>2. Anvil displacement<br>3. Hammer displacement<br>4. Full displacement of both hammer and anvil | At the end of the sequence, did any displacement occur? If so, was the capuchin displaced just from the anvil or hammer or both? If displacement occurred, make a comment with the ID or age/sex of the displacing and displaced individuals.<br>E.g.: "LAR displaces JOE" or<br>"Subadult male displaces juvenile"<br>**Important**: only code no displacement if other individuals were present but there was no displacement. If no other capuchins are visible, then leave it blank at "none". **Note**: only code displacement once. For example, if a juvenile is using tools and gets displaced, then code displacement for the juvenile's tool use sequence, but not for the sequence then started by the displacer. |
| | social attention | 1. no social attention<br>2. social attention | At the end of the sequence, did any other individuals pay attention to processing by the tool-user? Attention means peering at the tool-user while they are processing (does not have to include scrounging).<br>**Important**: if there was no opportunity for social attention (i.e., no other individuals around) then leave it blank at "none" and do not code "no social attention". |
| hammer stone | | | *To collect information about the hammerstones* |
| | hammer location start | 1. In hand<br>2. On anvil<br>3. Off anvil within reach<br>4. Off anvil walk<br>5. Carry in | The location of the hammerstone first used for the tool use sequence relative to the experimental anvil. Code the location where the hammerstone is at the beginning of the sequence. "In hand" means the capuchin is already holding it when the video starts, while "on anvil" means it is lying on the anvil and off anvil can be either "within reach" (within 1 body length of the anvil) or at a "walking distance" (>1 body length). Lastly the hammerstone can be "carried in" from out of view. |
| | hammer location end | 1. on anvil<br>2. off anvil within reach<br>3. off anvil out of reach<br>4. carry out | Code the hammerstone location again at the end of the sequence. Now the location can only be on the anvil, off anvil within reach (within 1 body length of the anvil), off anvil at a further distance or carry out if they take it. |
| | hammer ID | Enter the hammerstone ID as a comment, or if it's not marked/unidentifiable:<br>7. Unmarked<br>8. Unknown | Comment the ID of the hammerstone first used for the sequence, which can be one of the marked hammerstones or in the case of an unmarked hammerstone code "unmarked" but describe its appearance in the comments. |

#### 2.2 Coding of sequences with capuchins present

Table S3: Ethogram of behaviours coded per sequence with other capuchins present.

| behaviour | Modifiers | Modifier Options | Definition |
|---|---|---|---|
| Present | | | *Code once for each non-tool-using individual present during the tool use sequence. Make sure to assign age-sex class and possibly ID.* ***Note:*** *always code for every capuchin, also if individual also scrounges or does something else in sequence* |
| Social attention | | | *Sustained (>3 sec) attention to the tool user while they are on the anvil from a close distance (<2 meters). Only code once for each individual who does it.* |
| Displace | | | *Displace the tool user from the anvil. Code once for each individual who does it.* |
| Scrounge | | | *Scrounging is eating the items opened by the tool user during the sequence* |
| | Toleration | 1. Tolerated<br>2. Afterwards | Scrounging can be tolerated (occurring during the tool use, with the tool user still present and allowing it) or it can happen after the tool user has left the anvil.<br>**Note:** In the rare occasion that scrounging occurs despite aggression from the tool user, make a comment with ”stealing”. |
| Avoid | | | *Move away from the tool user* |
| | Aggression | 1. Aggression<br>2. No aggression | Whether the individual who avoids the tool user receives aggression from the tool user before moving away |

### 3 Supplemental results

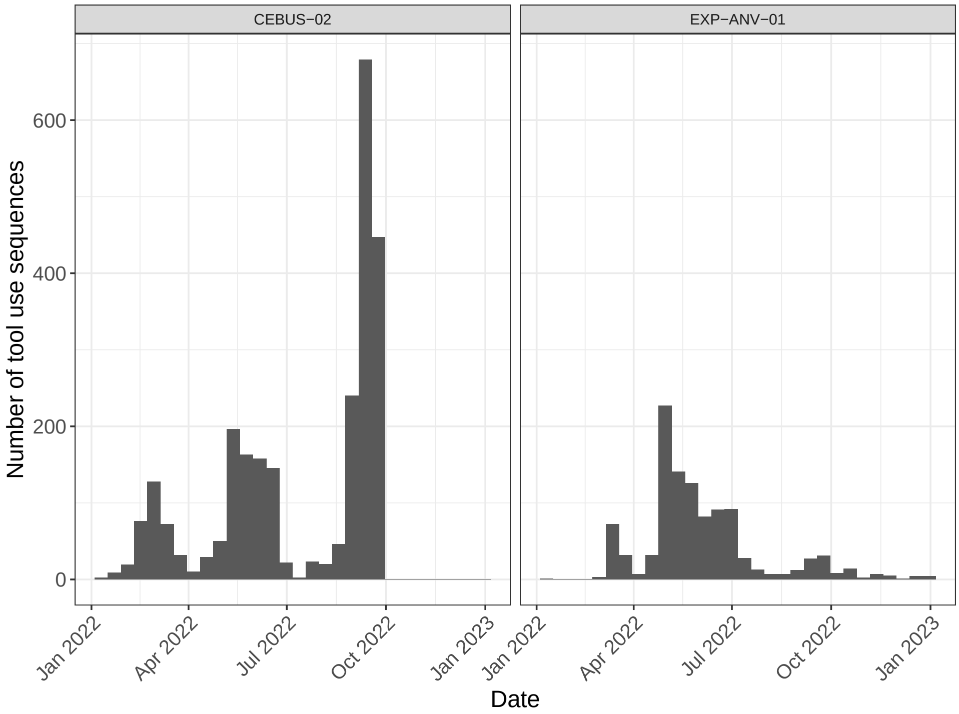


Figure S1: Number of tool use sequences observed per experimental anvil. At CEBUS-02, cameras ran 2022-01-10 until 2022-06-29 and 2022-07-16 until 2022-09-30. At EXP-ANV-01, cameras ran 2022-01-10 until 2022-07-16 and 2022-07-17 until 2023-01-01.

#### 3.1 Tool using proficiency

##### 3.1.1 Models estimating tool use efficiency

Table S4: Posterior mean model estimates of Model e1, a Gamma GLMM (Bayes $R^2$ = 0.24) examining factors influencing duration of tool use sequences. All estimated effects are on the log scale. The reference categories (the intercept) are juveniles (*age*), brown sea almonds (*item*), and stone (*anviltype*).

| | *Estimate* | *Est. Error* | *CI_95_low* | *CI_95_high* |
|---|---|---|---|---|
| *Multilevel Hyperparameters* | | | | |
| subjectID (11 levels) | 0.32 | 0.12 | 0.16 | 0.60 |
| *Regression Coefficients* | | | | |
| Intercept | 2.97 | 0.15 | 2.67 | 3.26 |
| Age (Subadult) | -0.69 | 0.24 | -1.17 | -0.24 |
| Age (Adult) | -0.55 | 0.25 | -1.03 | -0.06 |
| item (green sea almond) | 0.28 | 0.03 | 0.23 | 0.34 |
| item (red sea almond) | 0.40 | 0.07 | 0.28 | 0.52 |
| item (unknown sea almond) | 0.07 | 0.03 | 0.02 | 0.12 |
| anviltype (wood) | 0.01 | 0.02 | -0.04 | 0.05 |
| *Further Parameters* | | | | |
| shape | 4.02 | 0.11 | 3.82 | 4.23 |

Table S5: Posterior mean model estimates of Model e2, a Poisson GLMM (Bayes $R^2$ = 0.14) examining factors influencing number of pounds needed to open an item. All estimated effects are on the log scale. The reference categories (the intercept) are juveniles (*age*), brown sea almonds (*item*), and stone (*anviltype*).

| | *Estimate* | *Est. Error* | *CI_95_low* | *CI_95_high* |
|---|---|---|---|---|
| *Multilevel Hyperparameters* | | | | |
| subjectID (11 levels) | 0.17 | 0.07 | 0.08 | 0.36 |
| *Regression Coefficients* | | | | |
| Intercept | 1.67 | 0.09 | 1.50 | 1.85 |
| Age (Subadult) | -0.36 | 0.14 | -0.65 | -0.09 |
| Age (Adult) | -0.32 | 0.14 | -0.61 | -0.05 |
| item (green sea almond) | 0.24 | 0.03 | 0.19 | 0.30 |
| item (red sea almond) | 0.28 | 0.06 | 0.17 | 0.40 |
| item (unknown sea almond) | 0.07 | 0.03 | 0.01 | 0.12 |
| anviltype (wood) | -0.11 | 0.02 | -0.15 | -0.07 |

Table S6: Posterior mean model estimates of Model e2b, a Poisson GLMM (Bayes $R^2$ = 0.67) examining factors affecting the rate of pounding in tool use sequences. All estimated effects are on the log scale. The reference categories (the intercept) are juveniles (*age*), brown sea almonds (*item*), and stone (*anviltype*).

| | *Estimate* | *Est. Error* | *CI_95_low* | *CI_95_high* |
|---|---|---|---|---|
| *Multilevel Hyperparameters* | | | | |
| subjectID (11 levels) | 0.14 | 0.06 | 0.07 | 0.29 |
| *Regression Coefficients* | | | | |
| Intercept | -1.31 | 0.08 | -1.45 | -1.15 |
| Age (Subadult) | 0.30 | 0.12 | 0.07 | 0.56 |
| Age (Adult) | 0.29 | 0.13 | 0.01 | 0.52 |
| item (green sea almond) | -0.04 | 0.03 | -0.09 | 0.01 |
| item (red sea almond) | -0.10 | 0.06 | -0.22 | 0.02 |
| item (unknown sea almond) | 0.00 | 0.03 | -0.05 | 0.05 |
| anviltype (wood) | -0.12 | 0.02 | -0.16 | -0.07 |

##### 3.1.2 Models examining repositioning and peeling during tool use

Table S7: Posterior mean model estimates of Model e3a, a Poisson GLMM (Bayes $R^2$ = 0.17), examining factors influencing number of repositions during tool use sequences. All estimated effects are on the log scale. The reference categories (the intercept) are juveniles (*age*), brown sea almonds (*item*), and stone (*anviltype*).

| | *Estimate* | *Est. Error* | *CI_95_low* | *CI_95_high* |
|---|---|---|---|---|
| *Multilevel Hyperparameters* | | | | |
| subjectID (11 levels) | 0.55 | 0.20 | 0.28 | 1.04 |
| *Regression Coefficients* | | | | |
| Intercept | 0.07 | 0.25 | -0.48 | 0.53 |
| Age (Subadult) | -1.21 | 0.40 | -1.90 | -0.32 |
| Age (Adult) | -1.19 | 0.40 | -1.93 | -0.34 |
| item (green sea almond) | 0.21 | 0.08 | 0.04 | 0.37 |
| item (red sea almond) | 0.17 | 0.20 | -0.24 | 0.53 |
| item (unknown sea almond) | -0.05 | 0.08 | -0.20 | 0.10 |
| anviltype (wood) | 0.08 | 0.06 | -0.05 | 0.20 |

Table S8: Posterior mean model estimates of Model e3b, a Poisson GLMM (Bayes $R^2$ = 0.12) examining factors influencing number of peeling of an item during tool use sequences. All estimated effects are on the log scale. The reference categories (the intercept) are juveniles (*age*), brown sea almonds (*item*), and stone (*anviltype*).

| | *Estimate* | *Est. Error* | *CI_95_low* | *CI_95_high* |
|---|---|---|---|---|
| *Multilevel Hyperparameters* | | | | |
| subjectID (11 levels) | 0.25 | 0.21 | 0.01 | 0.82 |
| *Regression Coefficients* | | | | |
| Intercept | -1.19 | 0.28 | -1.76 | -0.67 |
| Age (Subadult) | -1.43 | 0.33 | -2.00 | -0.66 |
| Age (Adult) | -1.65 | 0.39 | -2.38 | -0.82 |
| item (green sea almond) | 1.29 | 0.20 | 0.90 | 1.68 |
| item (red sea almond) | 0.84 | 0.42 | 0.01 | 1.62 |
| item (unknown sea almond) | 0.21 | 0.25 | -0.29 | 0.69 |
| anviltype (wood) | -1.21 | 0.19 | -1.58 | -0.84 |

##### 3.1.3 Models examining mistakes during tool use

Table S9: Posterior mean model estimates of Model e4a, a zero-inflated Poisson GLMM (Bayes $R^2 = 0.18$) examining factors influencing number of misstrikes in a tool use sequence. All estimated effects are on the log scale. The reference categories (the intercept) are juveniles (*age*), brown sea almonds (*item*), and stone (*anviltype*).

| | *Estimate* | *Est. Error* | *CI_95_low* | *CI_95_high* |
|---|---|---|---|---|
| *Multilevel Hyperparameters* | | | | |
| subjectID (11 levels) | 1.84 | 0.52 | 1.05 | 3.03 |
| *Regression Coefficients* | | | | |
| Intercept | -2.31 | 0.73 | -3.84 | -0.98 |
| Age (Subadult) | -1.18 | 0.86 | -2.86 | 0.53 |
| Age (Adult) | -0.60 | 0.85 | -2.26 | 1.04 |
| item (green sea almond) | -0.71 | 0.50 | -1.77 | 0.19 |
| item (red sea almond) | 0.33 | 0.65 | -1.05 | 1.54 |
| item (unknown sea almond) | -0.10 | 0.30 | -0.70 | 0.46 |
| anviltype (wood) | -0.27 | 0.23 | -0.74 | 0.18 |
| *Further Parameters* | | | | |
| zi | 0.44 | 0.10 | 0.22 | 0.61 |

Table S10: Posterior mean model estimates of Model e4b, a zero-inflated Poisson GLMM (Bayes $R^2 = 0.10$) examining factors influencing number of item flying in a tool use sequence. All estimated effects are on the log scale. The reference categories (the intercept) are juveniles (*age*), brown sea almonds (*item*), and stone (*anviltype*).

| | *Estimate* | *Est. Error* | *CI_95_low* | *CI_95_high* |
|---|---|---|---|---|
| *Multilevel Hyperparameters* | | | | |
| subjectID (11 levels) | 0.21 | 0.18 | 0.01 | 0.69 |
| *Regression Coefficients* | | | | |
| Intercept | -1.14 | 0.27 | -1.70 | -0.66 |
| Age (Subadult) | -1.46 | 0.29 | -1.96 | -0.83 |
| Age (Adult) | -1.61 | 0.34 | -2.24 | -0.92 |
| item (green sea almond) | 1.25 | 0.19 | 0.87 | 1.63 |
| item (red sea almond) | 0.90 | 0.40 | 0.10 | 1.65 |
| item (unknown sea almond) | 0.15 | 0.24 | -0.34 | 0.62 |
| anviltype (wood) | -1.23 | 0.18 | -1.58 | -0.88 |
| *Further Parameters* | | | | |
| zi | 0.43 | 0.11 | 0.19 | 0.61 |

#### 3.2 Tool using development

Table S11: Posterior mean model estimates of Model dev1, a Poisson GLM (Bayes $R^2 = 0.20$) examining individual development of tool use proficiency over time by considering the number of pounds used to open a brown sea almond. All estimated effects are on the log scale and the reference individual is SMG, the oldest male in the sample.

| | *Estimate* | *Est. Error* | *CI_95_low* | *CI_95_high* |
|---|---|---|---|---|
| Intercept | 1.22 | 0.04 | 1.15 | 1.29 |
| days standardized | 0.09 | 0.04 | 0.01 | 0.16 |
| subjectID (PEA) | 0.23 | 0.06 | 0.13 | 0.35 |
| subjectID (BAL) | 0.34 | 0.05 | 0.24 | 0.43 |
| subjectID (TER) | 0.76 | 0.05 | 0.65 | 0.86 |
| subjectID (MIC) | 0.00 | 0.05 | -0.10 | 0.11 |
| subjectID (LAR) | -0.06 | 0.05 | -0.14 | 0.03 |
| subjectID (SPT) | -0.01 | 0.05 | -0.10 | 0.09 |
| subjectID (TOM) | 0.09 | 0.08 | -0.07 | 0.25 |
| subjectID (ZIM) | 0.54 | 0.09 | 0.36 | 0.71 |
| log(time) * subjectID (PEA) | -0.07 | 0.05 | -0.18 | 0.04 |
| log(time) * subjectID (BAL) | -0.09 | 0.05 | -0.19 | 0.32 |
| log(time)* subjectID (TER) | -0.17 | 0.05 | -0.28 | 0.04 |
| log(time) * subjectID (MIC) | -0.09 | 0.05 | -0.20 | 0.00 |
| log(time) * subjectID (LAR) | -0.03 | 0.05 | -0.13 | -0.06 |
| log(time) * subjectID (SPT) | -0.10 | 0.05 | -0.19 | -0.01 |
| log(time) * subjectID (TOM) | -0.06 | 0.05 | -0.21 | 0.08 |
| log(time) * subjectID (ZIM) | 0.16 | 0.08 | -0.01 | 0.32 |

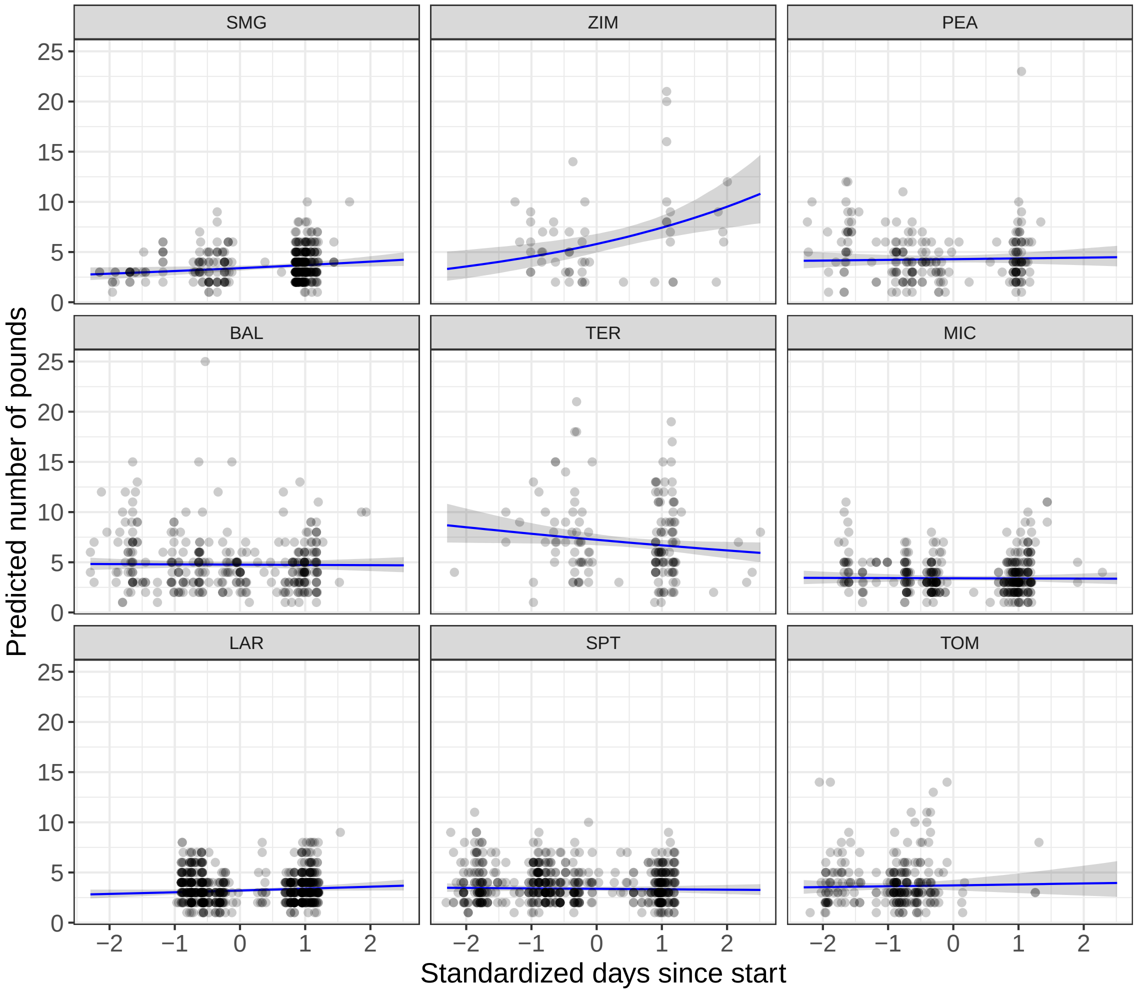


Figure S2: Change in the number of pounds over time for each individual tool user estimated by model dev1. Lines show posterior mean estimates from the Poisson GLM; shaded areas indicate 95% credible intervals. Points represent observed values from camera-trap data.

##### 3.2.1 Models examining social attention during tool use

Table S12: Posterior mean model estimates of Model socatt1, a Bernoulli GLMM (Bayes $R^2 = 0.26$) examining factors influencing the likelihood of an individual paying social attention to a tool use sequence. all estimated effects are on the logit scale. The reference categories (the intercept) are juveniles (*tooluser_age*), juveniles(*observer_agesex*), and CEBUS-02 (*location*).

| | *Estimate* | *Est. Error* | *CI_95_low* | *CI_95_high* |
|---|---|---|---|---|
| *Multilevel Hyperparameters* | | | | |
| sequenceID (907 levels) | 0.74 | 0.28 | 0.11 | 1.26 |
| *Regression Coefficients* | | | | |
| Intercept | -4.83 | 0.31 | -5.45 | -4.26 |
| Tool user (Subadult) | 1.10 | 0.24 | 0.64 | 1.58 |
| Tool user (Adult) | 0.84 | 0.33 | 0.18 | 1.46 |
| Observer (Subadult male) | -1.24 | 0.28 | -1.83 | -0.71 |
| Observer (Adult male) | -2.28 | 0.49 | -3.29 | -1.38 |
| Observer (Adult female) | -1.90 | 0.45 | -2.82 | -1.07 |
| location (EXP-ANV-01) | 0.86 | 0.22 | 0.43 | 1.30 |
| Capuchins present | -0.39 | 0.09 | -0.56 | -0.22 |
| Capuchins scrounging | 1.31 | 0.18 | 0.98 | 1.67 |

Table S13: Posterior mean model estimates of Model socatt1b, a Bernoulli GLMM (Bayes $R^2 = 0.08$) examining factors influencing the likelihood of an individual paying social attention to a tool use sequence. all estimated effects are on the logit scale. The reference categories (the intercept) are juveniles (*tooluser_age*), juveniles(*observer_agesex*), and CEBUS-02 (*location*).

| | *Estimate* | *Est. Error* | *CI_95_low* | *CI_95_high* |
|---|---|---|---|---|
| *Multilevel Hyperparameters* | | | | |
| tooluserID (10 levels) | 0.18 | 0.16 | 0.01 | 0.60 |
| *Regression Coefficients* | | | | |
| Intercept | -4.21 | 0.33 | -4.87 | -3.58 |
| Tool user (Subadult) | 0.96 | 0.29 | 0.43 | 1.56 |
| Tool user (Adult) | 0.52 | 0.36 | -0.18 | 1.24 |
| Observer (Subadult male) | -0.06 | 0.36 | -0.15 | 0.02 |
| Observer (Adult male) | -1.11 | 0.25 | -1.61 | -0.64 |
| Observer (Adult female) | -2.23 | 0.50 | -3.29 | -1.34 |
| Number of pounds | -1.76 | 0.43 | -2.65 | -0.97 |
| Number of mistakes | -0.07 | 0.21 | -0.49 | 0.33 |

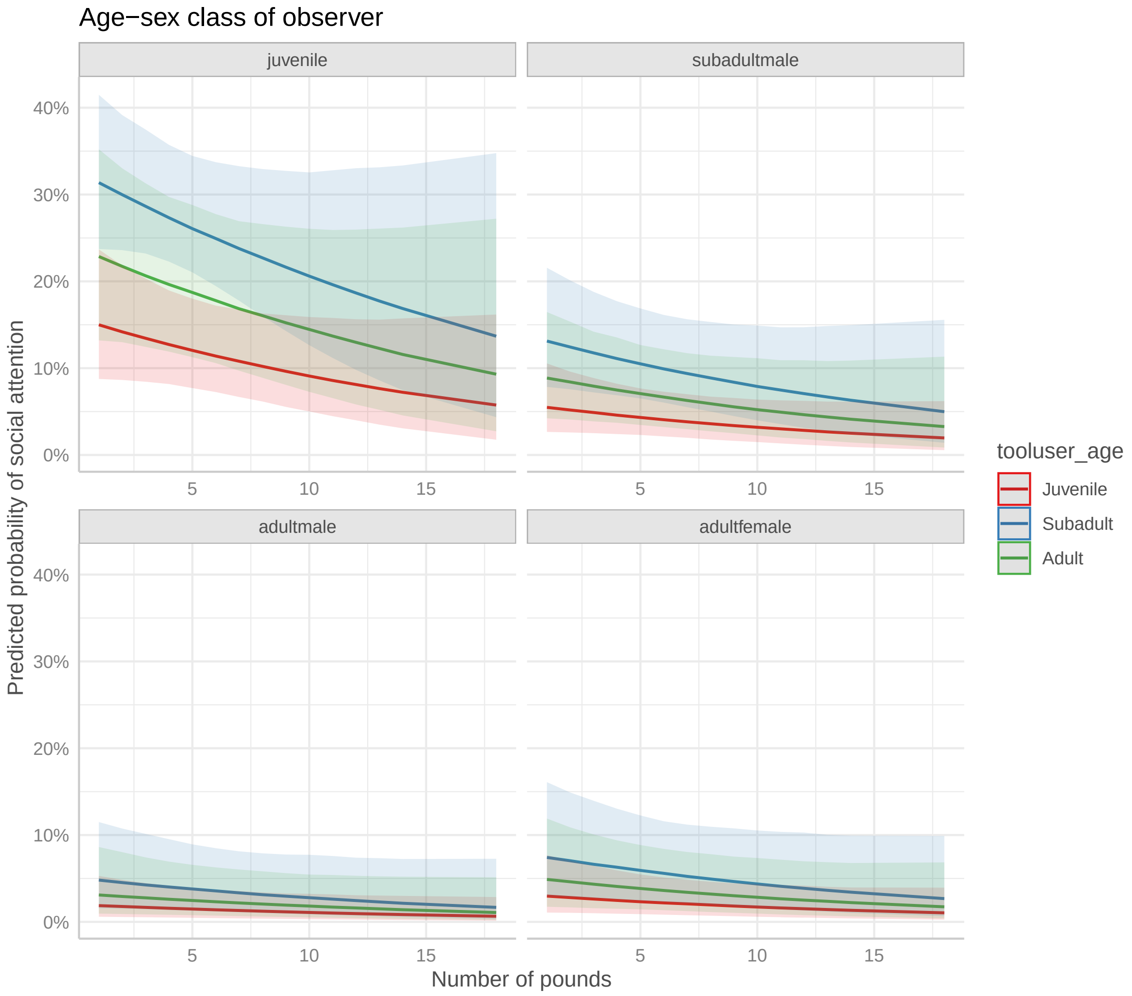


Figure S3: Predicted probabilities from model socatt1b of probabilities of social attention occurring, depending on the number of pounds in the sequence. Each facet represents an observer age-sex class, and the colour of the line the age of the tool user.
